# Parallel frequency-multiplexed aberration measurement for widefield fluorescence microscopy

**DOI:** 10.1101/2025.10.11.681535

**Authors:** Hyeonggeon Kim, Iksung Kang, Ryan Natan, Na Ji

## Abstract

Widefield fluorescence microscopy is widely used for imaging at subcellular resolution, but its performance in complex samples is degraded by optical aberrations. Because aberrations can vary spatially across the field of view (FOV), accurate aberration measurement and correction at multiple FOV locations are essential for achieving high-quality imaging over large areas. Here, we introduce parallel frequency-multiplexed aberration measurement (PFAM) to perform massively parallel aberration measurements across an extended FOV. We validated PFAM using fluorescent beads and demonstrated simultaneous measurement and effective correction of spatially varying aberrations at 125 FOV locations. To address the challenges of wavefront sensing in complex samples, we further developed PFAM-SIFT by integrating structured illumination, thereby achieving robust aberration measurement in both brain slices and the mouse brain in vivo. Together, PFAM and PFAM-SIFT provide accurate and scalable wavefront sensing solutions for widefield imaging, enabling simultaneous aberration measurement of spatially varying aberrations in complex biological samples.

## Introduction

Optical microscopy is an indispensable tool in biological imaging due to its noninvasive nature and capability for subcellular resolution. Among various microscopy modalities, standard widefield fluorescence microscopy—where the sample is uniformly illuminated and fluorescence signals from the entire FOV are simultaneously recorded by a camera—stands out for its high imaging speed, simplicity, and cost-effectiveness. However, like all optical imaging techniques, widefield fluorescence microscopy suffers from resolution degradation and contrast reduction caused by optical aberrations. These aberrations arise from imperfections within the imaging system (system aberrations), as well as from refractive index heterogeneities within biological tissues and/or refractive index mismatches between the sample and the immersion medium of the microscope objective (sample-induced aberrations).

Adaptive optics (AO) can restore diffraction-limited imaging performance by accurately measuring and correcting these aberrations^1–3^. AO typically involves two main steps: wavefront sensing, which measures the aberrated wavefront, and wavefront correction, where a wavefront conjugate (of the opposite phase) to the measured aberration is applied using a correction device such as a liquid-crystal spatial light modulator (SLM) or a deformable mirror (DM). Alternatively, image degradation caused by aberrations can be corrected computationally through methods such as deconvolution^4–6^ or machine learning algorithms^7–9^.

Several AO methods, each employing distinct strategies for wavefront sensing, have previously been developed for widefield fluorescence microscopy. Their wavefront sensing methods broadly fall into two categories: direct wavefront sensing (DWS) and indirect wavefront sensing. DWS uses a dedicated wavefront sensor, often a Shack– Hartmann sensor^10^, to directly measure the local phase gradient of an aberrated wavefront. This usually requires fluorescence signal from a three-dimensionally (3D) confined guide star, such as an exogenously introduced fluorescent bead^11,12^, or a fluorescent volume generated by confocal^13^ or two-photon excitation^14,15^. In contrast, indirect wavefront sensing methods (also referred to as sensorless AO) do not rely on external sensors. Instead, these methods infer aberrations either by measuring image displacements across different pupil zones^16^ or by measuring how image-quality metrics (e.g., spatial frequency content^13,17,18^) vary with known aberrations applied to the wavefront correction device. Additionally, computational approaches, such as phase retrieval algorithms, reconstruct phase information from intensity measurements of the three-dimensional point spread function (PSF) using sub-diffraction fluorescent objects^19–21^.

In complex samples such as biological tissues, wavefront distortion accumulates as light propagates through an optically inhomogeneous sample volume. As a result, optical aberrations can vary spatially and the area within which aberrations are similar can be smaller than the image FOV. Correction based on the aberration measured at one location within the FOV can lead to wavefront errors in other locations, limiting the effective correction area and diminishing the overall performance of widefield microscopy.

For AO to improve image quality over the entire FOV in widefield microscopy, aberrations at different FOV locations need to be measured and corrected. However, conventional AO methods, which either measure local aberrations at a single location or average them across the FOV, are inherently limited in their capacity to provide optimal aberration correction across large FOVs. Methods capable of accurately measuring and correcting aberrations across multiple FOV locations are needed for achieving full-field AO correction.

Several AO approaches have been proposed to address this challenge. Conjugate AO, for example, extends the corrected FOV in the presence of spatially varying aberrations by conjugating wavefront correction devices to specific aberrating layers^22–24^. Although effective for aberrations induced by a few dominant layers, this method is not suitable for general cases where aberrations arise from a continuous volume, as is typical for biological tissues. A pupil-segmentation approach measures field-dependent aberrations from local shifts in images that were formed by fluorescence emitted through different pupil segments^25,26^. However, the reduction of emission NA during aberration measurement limits its application to thin samples or requiring light-sheet excitation. Another method places a random phase mask in the fluorescence emission path to generate a speckle pattern on camera and, using fluorescence from multiple guide stars, simultaneously measures the aberrations at guide star locations from local speckle shifts^27^. However, because speckles spread energy over many pixels, achieving high multiplexing capability becomes challenging under low-light conditions. Therefore, achieving both high accuracy and high throughput in parallel aberration measurements from complex samples remains a significant challenge.

In this paper, we introduce a Parallel Frequency-multiplexed Aberration Measurement (PFAM) approach that simultaneously measures aberrations at multiple locations across the FOV of a widefield fluorescence microscope. We first validate the method by performing massively parallel aberration measurements on fluorescent beads with spatially varying aberrations. The measured aberrations are then used to digitally correct their widefield images via patch-wise non-blind Richardson–Lucy deconvolution (RLD)^28,29^, improving image quality over the entire FOV.

We further broaden the applicability of PFAM to complex, extended samples with PFAM-SIFT, which integrates PFAM with structured illumination (SI) and Fourier transform (FT). Creating additional high spatial frequencies for aberration measurement and isolating in-focus signals from out-of-focus background, SI allows PFAM-SIFT to maintain robust wavefront sensing performance regardless of the sample’s lateral or axial dimension, as demonstrated by a variety of samples including large fluorescent beads, fluorescent films, and mouse brain slices, including for spatially varying aberrations. Finally, we apply the method to live mouse brain imaging and show that PFAM-SIFT reliably measures sample-induced aberrations under in vivo condition.

## Results

### Frequency-multiplexed wavefront sensing on widefield fluorescence microscopy

We implemented PFAM in widefield fluorescence microscopy by placing a DM in the detection path to measure and, if desired, correct aberrations of the emitted fluorescence (**Fig. 1a**). The DM had 169 mirror segments (gray segments, left inset, **Fig. 1a**) with individual control for each segment’s tip, tilt, and piston values. Placed in conjugation to the pupil planes of the microscope objective and the camera focusing lens, it modifies the wavefront of the fluorescence reflected off it before image formation on the camera.

**Figure 1.**
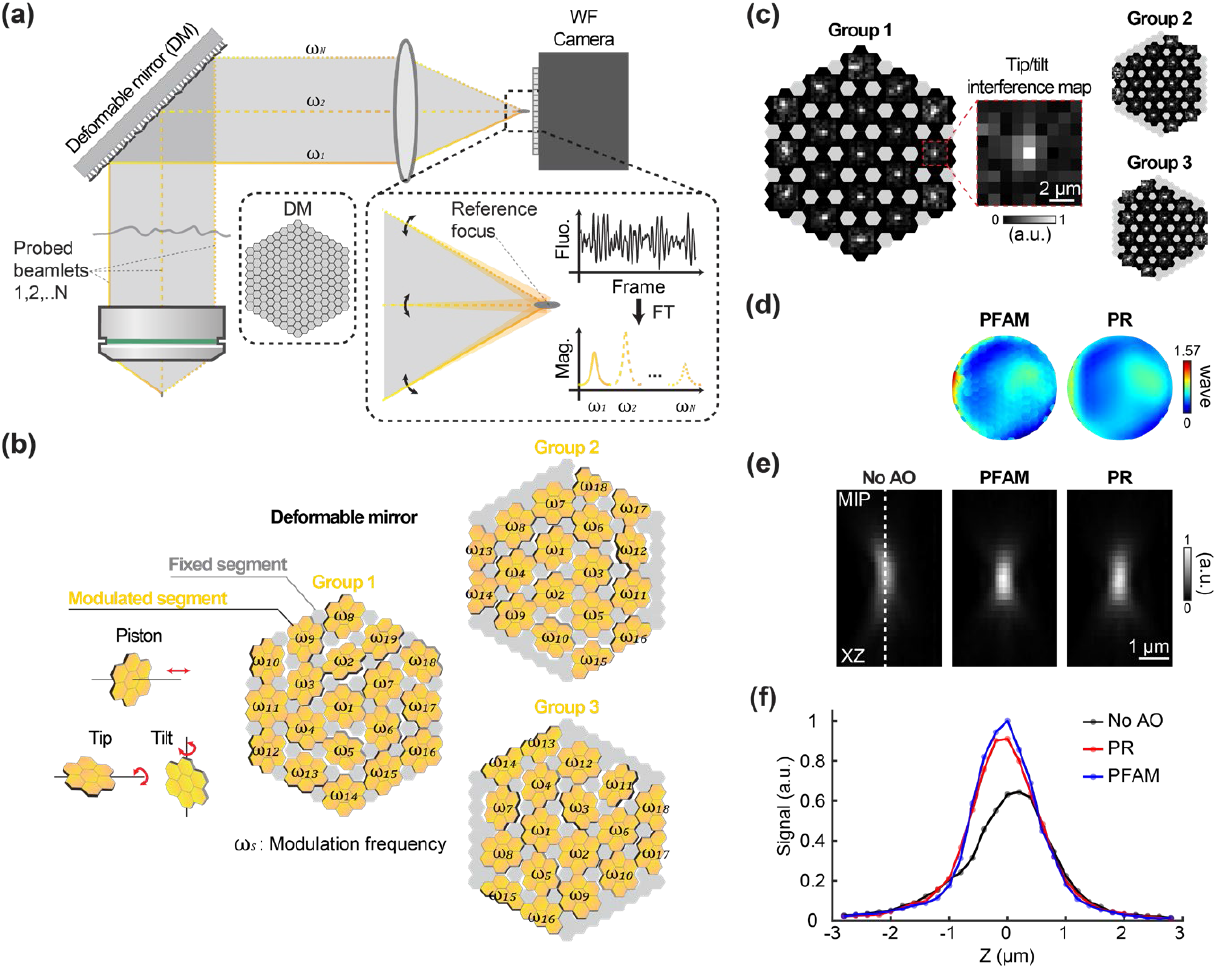
Principle and implementation of parallel frequency-multiplexed aberration measurement (PFAM) for widefield fluorescence microscopy. (**a**) Schematic of PFAM implemented in a widefield fluorescence microscope. A deformable mirror (DM, inset) is placed at a plane conjugate to the objective’s back pupil for both aberration measurement and correction. Inset: (Left) Enlarged view of the image plane at the widefield (WF) camera, showing *N* probe beamlets (gold) scanned around a stationary reference focus (gray) by controlling tip and tilt of their corresponding DM macro-segments; (Upper right) At each tip/tilt configuration, phase of probe beamlet is modulated at a unique frequency *ωs* (*ω*_1_, *ω*_2_, …, *ω*_*N*_) by controlling the piston of its corresponding macro-segment, modulating the fluorescence signal resulting from the interference between the probe beamlet and the reference focus at frequency *ω*_*s*_; (Lower right) Interference strength between each probe beamlet and the reference focus is extracted by performing a Fourier transform (FT) on the fluorescence signal trace and identifying the Fourier magnitude at *ω*_*s*_. (**b**) DM grouping configuration. Modulated regions (gold) consist of macro-segments formed by grouping adjacent DM segments with piston, tip, and tilt control. Remaining segments (gray) are held static to generate the reference focus. To increase sampling density of phase gradient measurement, three DM groups are typically used sequentially (see other groupings in **Supplementary Fig. 1**). (**c**) Tip/tilt interference maps for macro-segments across three DM groups, from which phase gradients are measured. Inset: Example interference map (scaled to the sample plane) of a macro-segment. (**d**) Corrective wavefronts for system aberration measured using PFAM from a 0.5-μm-diameter fluorescent bead (left) and phase retrieval from a 0.2-μm-diameter fluorescent bead (right). Both PFAM and PR were iterated twice. (**e**) Maximum intensity projection (MIP) images in XZ of a 0.5-μm-diameter fluorescent bead acquired without adaptive optics (AO, left), with corrective wavefront from PFAM (middle), and with corrective wavefront from phase retrieval (right). (**f**) Axial fluorescence intensity profiles along dashed lines in (**e**).

PFAM measures the phase gradients and offsets that must be applied to the fluorescence wavefront emitted by a fluorophore so that their light rays intersect at the same point on the camera with a common phase, enabling maximum constructive interference and generating an ideal, aberration-free image of the fluorophore (**Supplementary Fig. 1**, Methods). During PFAM measurement, we group multiple mirror segments into a macro-segment with a planar surface and vary the tip, tilt, and piston of these macro-segments (gold segments, Group 1, **Fig. 1b**) while keeping the rest of mirror segments stationary (gray segments, Group 1, **Fig. 1b**).

We first measure the phase gradient that needs to be applied to each macro-segment for the beamlets reflected off them to converge at the same point on the camera. By varying the tip and tilt angles of macro-segments, we scan probe beamlets (represented by gold lines, **Fig. 1a**) reflected off these macro-segments around the reference focus (gray focus, right inset, **Fig. 1a**) formed by beamlets (gray beam, **Fig. 1a**) that reflect off fixed segments (gray segments, Group 1, **Fig. 1b**). The tip/tilt angles of each macro-segment are chosen randomly from an array of angles that scan the probe beamlets over ±318 μm in two dimensions (2D) around their original location in the camera plane (corresponding to ±4.2 μm in the sample plane) (Step 1, **Supplementary Fig. 1**).

At each set of tip and tilt angles, we modulate the phase offset of each probe beamlet at a distinct frequency ω_*s*_ (*s* = 1, 2, …, *N*; *N*: the total number of macro-segments) by varying the piston value of the corresponding macro-segment and record a camera frame for each phase offset value. We then select a FOV location (a single camera pixel or a group of pixels representing structures of interest) and extract its pixel value from each frame, acquiring a fluorescence trace (right inset, **Fig. 1a**; Step 2, **Supplementary Fig. 1**). We Fourier transform this trace and record the Fourier magnitude for each modulation frequency ω_*s*_ (right inset, **Fig. 1a**; Step 3, **Supplementary Fig. 1**).

If a probe beamlet does not overlap with the reference focus, the modulation on its phase would not affect the recorded fluorescence signal, leading to zero Fourier magnitude. If they do overlap, their interference and the modulation on probe beamlet phase would lead to changes in signal and nonzero Fourier magnitude at the modulation frequency, with maximal Fourier magnitude observed at the tip/tilt angles that lead to maximal overlap between the probe beamlet and the reference focus. Because each probe beamlet is tagged with a unique modulation frequency ω_*s*_, the contribution to the interference signal by all probe beamlets can be extracted simultaneously by reading out the Fourier magnitudes at ω_*s*_ (*s* = 1, 2, …, *N*) (red circles, Step 3, **Supplementary Fig. 1**).

Repeating this measurement across all tip/tilt angles (Step 4, **Supplementary Fig. 1**) generates a 2D Fourier magnitude map for each macro-segment (**Fig. 1c**, the zoomed-in inset shows the map for one example macro-segment; Step 5, **Supplementary Fig. 1**). We then compute the weighted centroid of each map to identify the tip/tilt angles that produce maximal Fourier magnitude, which inform on the phase gradient of the corresponding probe beamlet. When these tip/tilt angles are applied to the corresponding macro-segment, the macro-segment directs the reflected probe beamlet to maximally overlap with the reference focus.

In our implementation, most macro-segments included seven neighboring segments (**Fig. 1b**) sharing the same tip and tilt angles and forming a planar surface, to enhance the strength of the probe beamlet and its maximal Fourier magnitude. To densely sample the phase gradients across the wavefront, we repeated the above phase gradient measurement for different groups of modulated and fixed DM segments. For higher measurement accuracy, segments were divided into three groups (Group 1, 2, and 3, **Fig. 1b**), with a 2D Fourier magnitude map measured for each group (**Fig. 1c**; Step 6, **Supplementary Fig. 1**). Alternatively, for faster measurement, a coarser grouping into two or one group was used (Step 1, **Supplementary Fig. 1**). For segments that were included in multiple macro-segments in different groups, final phase gradient values were calculated by averaging the measurements across the different groups. With the phase gradients known, we compute the phase offset for each segment to constructively interfere at the focus using a zonal wavefront reconstruction method^30,31^, which gives rise to the final corrective wavefront (“PFAM”, **Fig. 1d**).

For large aberrations, the initial reference focus can be substantially distorted and enlarged; in such cases, the above process may be iterated 2 to 3 times to reach the optimal corrective wavefront.

PFAM is based on a frequency-multiplexed aberration measurement principle that we previously developed for measuring aberrations in point-scanning microscopy including confocal, two-photon, and three-photon fluorescence microscopy, as well as third-harmonic generation microscopy^32–35^. However, unlike these earlier methods that use frequency-multiplexing to correct the aberrations in the excitation light path, PFAM is implemented in the fluorescence emission path. The corrective wavefront can be applied to the DM to physically correct aberrations of the emission light. Alternatively, its conjugate wavefront describes the aberration experienced by the fluorescence and can be used to compute a PSF for image deconvolution. As demonstrated below, the parallel recording of fluorescence signal across the entire FOV in widefield microscopy enables us to simultaneously measure spatially varying aberrations across the FOV.

### Microscope setup and system aberration correction

The microscope setup included a widefield single-photon fluorescence light path and a two-photon excitation (TPE) light path (**Supplementary Fig. 2**, Methods). A 488-nm or 561-nm continuous-wave laser was used for widefield excitation. The excitation light reflected off a spatial light modulator (SLM; Forth Dimension Displays Ltd., QXGA-3DM) that was optically conjugated to the objective focal plane. The SLM had a flat pattern for standard widefield microscopy; when a structured illumination pattern at the objective focal plane was needed, the SLM displayed a hexagonal binary pattern. The TPE path was used to generate a two-photon-excited guide star for direct wavefront sensing (DWS), which provided ground-truth aberration measurement and correction to validate PFAM performance. Alternatively, ground-truth aberrations were acquired via phase retrieval with the Gerchberg–Saxton algorithm^36^ from 3D stacks of 0.2-µm-diameter fluorescent beads.

We first tested the performance of PFAM in correcting the system aberration at the center of the FOV of the widefield microscope. During PFAM, we acquired 2D images of 0.5-µm-diameter fluorescent beads at a camera frame rate of 115 Hz (**Supplementary Video 1**). Using the averaged pixel value over 5×5-pixel regions centering at five beads at the FOV center, PFAM returned a corrective wavefront (“PFAM”, **Fig. 1d**), which closely matched the corrective wavefront obtained by phase retrieval from a 3D stack of a 0.2-µm-diameter bead (“PR”, **Fig. 1d**; root-mean-square error (RMSE) between the two wavefronts: 0.086 waves). We acquired 3D image stacks of a 0.2-µm-diameter bead without aberration correction, with corrective wavefront from PFAM, or with corrective wavefront from phase retrieval. Comparing the axial images of the bead (maximum intensity projection (MIP) in the XZ plane, **Fig. 1e**), we found that PFAM and phase retrieval similarly improved axial resolution (**Fig. 1e**) and increased signal (**Fig. 1f**). In addition, PFAM with coarser grouping of DM segments (**Supplementary Fig. 1**) also improved axial resolution and increased signal to a similar extent (**Supplementary Fig. 3**).

### Validation with artificially introduced aberrations

We further validated PFAM’s accuracy by using it to correct aberrations artificially introduced onto the DM. Applying astigmatism, coma, and spherical aberration (∼0.36 waves RMS) to the DM reduced the peak signal of a 0.5-µm-diameter bead to 0.25×, 0.28×, and 0.32× of the aberration-free signal, respectively. After 1 iteration of PFAM, the corrective wavefront led to a 3.9×, 2.7×, and 2.5× signal improvement and reduced RMSE to 0.057, 0.112, and 0.1 waves (**Fig. 2a-c**). With three iterations, peak signal intensity recovered to near-ideal levels, and the residual wavefront RMSE was reduced to near the Rayleigh criterion of 0.07 waves (**Fig. 2d-f**).

**Figure 2.**
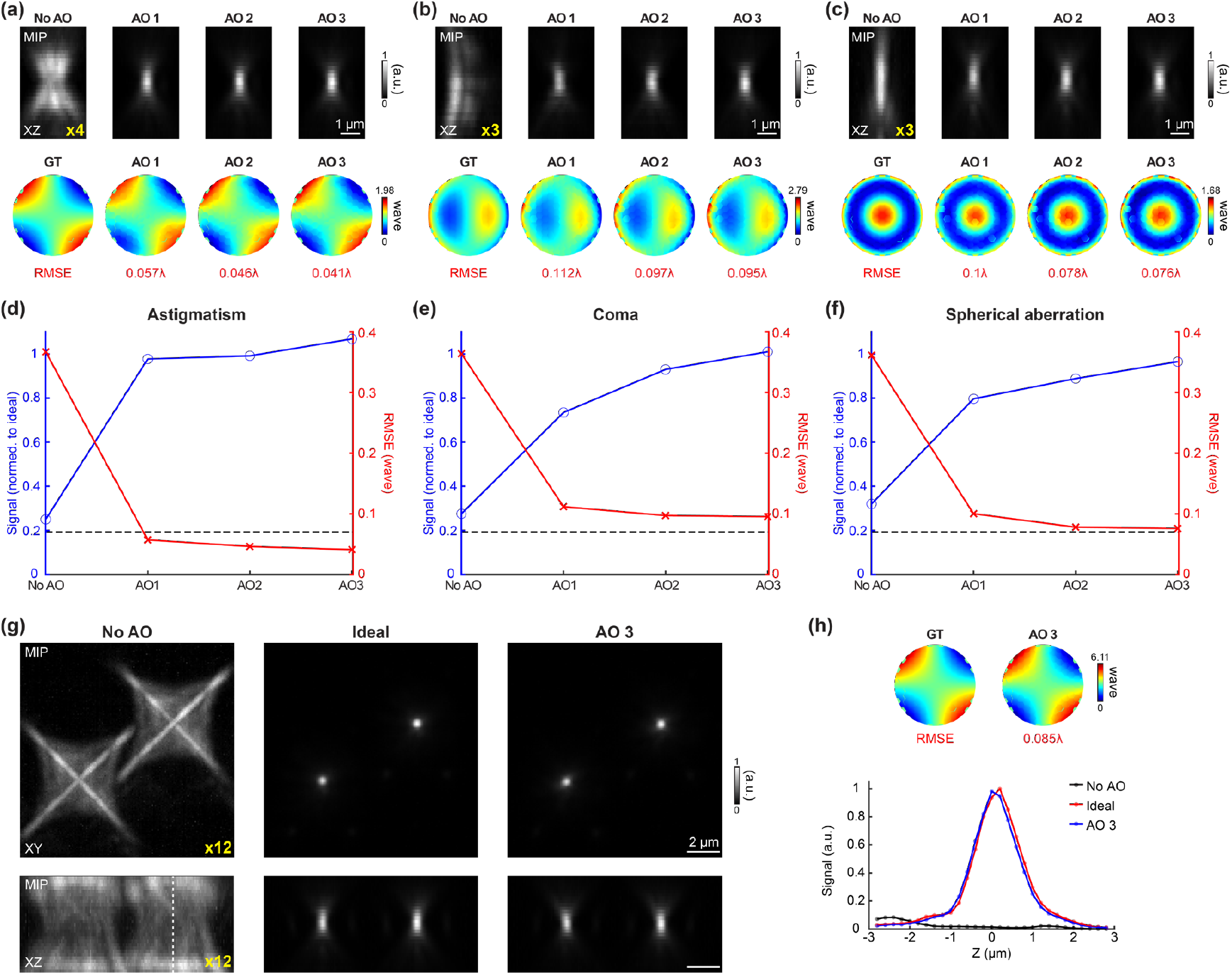
Correction of artificial aberrations using PFAM. Artificial aberrations applied to DM—astigmatism, coma, and spherical aberration of ∼0.36 waves RMS—were measured and corrected using PFAM over up to three iterations at a camera frame rate of 115 Hz. (**a-c**) (Top) Maximum intensity projections (MIPs) in XZ of a 0.5-μm-diamter fluorescent bead without aberration correction (“No AO”) and after one, two, or three PFAM iterations (“AO 1”, “AO 2”, “AO 3”). For visualization, pixel values in “No AO” images were artificially boosted, as indicated by yellow labels. (Bottom, from left to right) Ground-truth (GT) corrective wavefront (i.e., the opposite of aberration applied to DM) and corrective wavefronts measured by one, two, or three iterations of PFAM; RMSE: RMS errors between GT and measured wavefronts. (**d-f**) Peak signal (blue) of a 0.5-μm-diamter bead and RMSE (red) for No AO and across three PFAM iterations for each artificial aberration. Black dashed lines: diffraction-limited resolution. (**g**) PFAM correction of a large astigmatism aberration of 1.10 waves RMS. MIPs in XY and XZ for “No AO,” “Ideal” (no artificial aberration), and “AO 3” (after three iterations of PFAM) conditions. (**h**) GT corrective wavefront and measured corrective wavefront with line intensity profiles along the dashed lines in (**g**).

PFAM can successfully correct stronger artificial aberrations. For example, a large astigmatism aberration (1.10 waves RMS; 6.11 waves peak-to-valley) was introduced to the DM, severely degrading resolution and reducing the peak signal of a 0.5-µm-diameter bead by 12× (“No AO” versus “Ideal”, **Fig. 2g**). After 3 iterations of PFAM, the correction wavefront fully recovered the resolution and brightness (“AO3”, **Fig. 2g; Fig. 2h**), leading to a residual wavefront RMSE of 0.085 waves.

### Massively parallel measurement of spatially varying aberrations

In widefield fluorescence microscopy, the sample is illuminated broadly, and a camera detects fluorescence emission in parallel from all fluorescent structures within the FOV (e.g., fluorophore 1 and 2, **Fig. 3a**). Fluorescence originating from different FOV positions may accumulate distinct aberrations and thus have distinct brightness variations and Fourier transform magnitudes during PFAM (inset, signal traces and their Fourier transforms at FOV locations 1 and 2, **Fig. 3a**), reflecting the different phase gradients required to make their beamlets intersect at the same pixels on camera.

**Figure 3.**
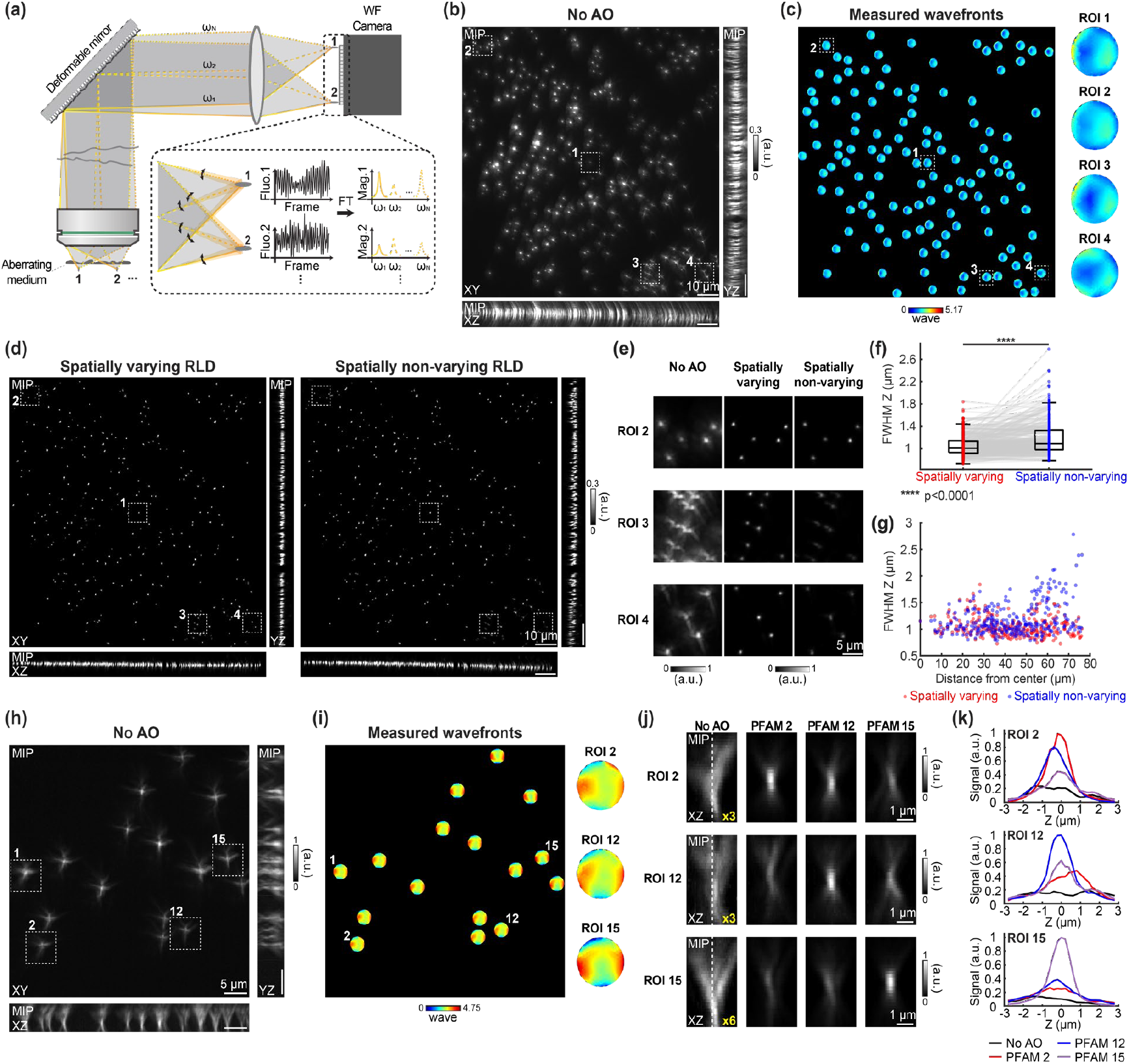
Massively parallel measurements of spatially varying aberrations using PFAM. (**a**) Schematics of simultaneous PFAM at multiple image locations enabled by parallel acquisition of fluorescence signals across the widefield (WF) camera. Inset: Fluorescence signals measured at distinct FOVs and their Fourier transform (FT) reflect different interference strengths between probe beamlets and reference focus at each FOV location due to spatially varying aberrations. (**b**) Maximum-intensity projections (MIPs) in XY, XZ, and YZ planes of 0.5-µm-diameter fluorescent beads beneath a thin aberrating layer that induces spatially varying aberrations over 120 µm × 120 µm (pixel size: 86 nm). Images are saturated for visibility. (**c**) Corrective wavefronts at 125 regions of interest (ROIs), measured by PFAM at 0.172 µm pixel size using 2×2 camera pixel binning readout at a camera frame rate of 77 Hz. Insets: Example wavefronts at ROIs 1-4 in (**b**). (**d**) XY, XZ, and YZ MIPs after Richardson–Lucy deconvolution (RLD) using spatially varying wavefronts (“spatially varying RLD”) or the corrective wavefront measured at ROI 1 (“spatially non-varying RLD”). (**e**) Magnified XY MIPs at ROIs 2–4 of “No AO” images, deconvolved images with spatially varying wavefronts, and deconvolved images with wavefront measured at ROI 1. Gray scale for ‘No AO’ image is normalized to its minimal and maximal pixel values, while deconvolved images share the same minimal and maximal values for gray scale. (**f**) Axial full widths at half maximum (FWHMs) for beads processed with spatially varying (red) and non-varying (blue) RLD. Box plots: center line, median; box edges, 25th/75th percentiles; whiskers, ±2.7×SD. (**g**) Axial FWHMs plotted as a function of bead distance from the FOV center (ROI 1) for beads processed with spatially varying (red) and non-varying (blue) RLD. (**h**) XY, XZ, and YZ MIPs of 0.5-µm-diameter fluorescent beads with spatially varying aberrations over 52 µm × 52 µm (pixel size: 86 nm). (**i**) Corrective wavefronts at 16 bead positions measured with PFAM at a camera frame rate of 50 Hz. Insets: Example corrective wavefronts at ROI 2, 12, and 15. (**j**) XZ MIPs of beads at ROIs 2, 12, and 15 acquired with No AO, corrective wavefronts acquired by PFAM at ROI 2, 12, and 15, respectively. (**k**) Axial intensity profiles along dashed lines in (**j**).

Because the camera records fluorescence across the FOV, PFAM can simultaneously measure aberrations at different locations: by performing Fourier transforms on the recorded signal traces of region of interests (ROIs) at different FOV positions, one can obtain location-specific tip/tilt interference maps and phase gradient measurements across the entire FOV.

To demonstrate its ability to perform massively parallel measurements of spatially varying aberrations, we used PFAM to measure the aberrations introduced by a thin layer of transparent nail polish above 0.5-µm-diameter fluorescent beads on a coverslip. Maximum intensity projections (MIPs) in the XY, XZ, and YZ planes (**Fig. 3b**), along with magnified XY views (“No AO”, **Fig. 3e**), revealed that bead images were distorted in a position-dependent manner by spatially varying aberrations.

We first performed one round of PFAM on a bead in the center of the FOV (ROI 1, **Fig. 3b**). Applying the corrective wavefront for this bead to the DM, we acquired 2D images of beads in a 120 μm × 120 μm FOV undergoing PFAM (**Supplementary Video 2**). Grouping nearby fluorescent beads into 125 ROIs (mean area: 34.10 μm^2^) and analyzing their signals, we measured spatially varying aberrations over the FOV in parallel, resulting in a total of 125 corrective wavefronts (**Fig. 3c, Supplementary Video 2**). Example corrective wavefronts from four ROIs (ROI 1-4, inset, **Fig. 3c**) clearly illustrate the spatial variability of the aberrations. To quantify this spatial variability, we calculated the RMS difference between each ROI’s wavefront and the wavefront measured at the FOV center (wavefront for ROI 1, inset, **Fig. 3c**). The differences ranged from 0.064 waves to 0.54 waves, with a mean of 0.19 waves and a standard deviation of 0.087 waves.

With the corrective wavefront for each ROI known, image degradation caused by spatially varying aberrations can be corrected computationally using non-blind deconvolution based on RLD. To evaluate the importance of using local aberration for deconvolution, we compared images deconvolved with ROI-specific wavefronts with images deconvolved with the wavefront measured from the central bead (ROI 1, **Fig. 3c**) applied uniformly across the FOV. MIP views in XY, XZ, and YZ planes show that spatially varying RLD effectively corrected field-dependent aberrations, aligning beads to a common Z plane with similar axial sizes (spatially varying RLD, **Fig. 3d**). In contrast, the spatially non-varying RLD led to artifactual Z displacements and varied axial sizes (spatially non-varying RLD, **Fig. 3d**). Enlarged XY views of ROIs 2–4 further highlight the superior improvement in the brightness and morphology of beads with ROI-specific corrections (**Fig. 3e**).

To quantify resolution enhancement, we measured the axial full width at half maximum (FWHM) of fluorescent beads across the FOV. Beads deconvolved using spatially varying wavefronts exhibited significantly narrower FWHMs compared to those deconvolved with a single, spatially invariant wavefront (**Fig. 3f**). When spatially varying wavefronts were used, the mean axial FWHM for these 0.5-µm-diameter beads was 1.05 μm with a standard deviation of 0.19 μm. In contrast, using a spatially invariant wavefront resulted in a larger mean FWHM of 1.19 μm and standard deviation of 0.31 μm (diffraction-limited axial resolution for NA-1.1 objective at 515 nm emission: 1.13 μm). This improvement became more pronounced with increasing distance from the FOV center (**Fig. 3g**), confirming that PFAM effectively extends high-resolution imaging across a large FOV.

We further validated the effectiveness of parallel aberration measurement by PFAM by optically correcting aberrations through direct application of the measured corrective wavefronts to the DM. As in **Fig. 3a**, spatially varying sample-induced aberrations were introduced by a thin nail-polish layer over sparsely distributed 0.5-µm-diameter fluorescent beads, which caused strong position-dependent distortions in the XY, XZ, and YZ MIP images (**Fig. 3h**). We first performed two PFAM iterations on the bead at ROI 1 to obtain a baseline corrective wavefront. Applying the corrective wavefront on the DM, PFAM then simultaneously measured the remaining spatially varying wavefronts at 16 bead positions for final corrective wavefronts (**Fig. 3i**), with representative wavefronts at ROIs 2, 12, and 15 clearly demonstrating spatial variability (inset, **Fig. 3i**).

To evaluate how the measured corrective wavefronts improved image quality, we applied the locally measured corrective wavefronts at ROI 2, 12, or 15 to the DM. As expected, the corrective wavefront acquired at each bead location consistently yielded the greatest improvement in the bead’s corresponding XZ MIP images (**Fig. 3j**) and axial signal profiles (**Fig. 3k**). As above, ROI-specific wavefronts can also be used for patch-wise non-blind deconvolution to computationally correct spatially varying aberrations (**Supplementary Fig. 4**).

In contrast to computational approaches^8,20^ that require the pixel size to satisfy the Nyquist sampling criterion, PFAM remains effective even when the pixel size is larger than the Nyquist limit. Given the lateral resolution of 0.234 μm for our microscope (Abbe’s criterion, 1.1 NA objective, 515 nm emission), Nyquist sampling requires a pixel size of 0.117 μm or smaller at the sample plane, which was satisfied by our pixel size of 0.086 μm. We performed 2×2, 4×4, or 8×8 pixel averaging on the images acquired during PFAM and used the resulting frames, which no longer satisfied Nyquist criterion, to measure astigmatism, coma, and spherical aberrations applied to the DM (**Supplementary Fig. 5a**). We found that PFAM remained effective across 2×2, 4×4, and 8×8 pixel averaging for all aberrations (**Supplementary Fig. 5b-d**) with only a slight increase in the RMS error of the measured corrective wavefronts at higher pixel averaging levels (**Supplementary Fig. 5e**). The fact that PFAM is highly robust to image undersampling enabled us to leverage camera pixel binning readout to reduce image acquisition time and computational load, in order to accelerate wavefront sensing across large FOVs. It also makes PFAM suitable for measuring aberrations in optical systems with large FOVs and pixel sizes that do not meet the Nyquist sampling criterion.

### Robust aberration measurement for spatially extended fluorescent structures using structured illumination

As implemented above, PFAM works well with fluorescent features (e.g., 0.5-µm-diameter beads) that are comparable in size to the reference focus (defined by diffraction limit at 1.1 NA and 515 nm emission: lateral 0.234 µm, axial 1.13 µm). This is because if there are few fluorophores outside the reference focus, when the probe beamlets do not intersect the reference focus, a substantial drop in brightness occurs for pixels within the reference focus. However, when fluorescent features extend substantially beyond the diffraction limit laterally or axially, the above brightness drop is compensated by beamlets that originate from fluorophores laterally or axially outside the diffraction limit and deviate from their own reference foci, reducing signal variation during phase gradient measurements. Widefield microscopy also captures fluorescence from out-of-focus features, which increases photon noise and further reduces the accuracy of PFAM.

Inspired by AO approaches developed for structured illumination microscopy (SIM)^18,37,38^, we integrated PFAM with structured illumination (SI) and Fourier transform (FT) and demonstrated that PFAM-SIFT can improve aberration measurement accuracy in both laterally and axially extended structures. In our implementation, we displayed a hexagonal binary pattern on the SLM, which generated six diffraction spots at the objective back pupil plane and a hexagonal lattice SI pattern at the objective focal plane (**Fig. 4a**). In the spatial frequency space, the SI corresponded to six spatial frequency components of magnitude 0.43k_0_ (k_0_: cutoff frequency defined by the objective NA) evenly spaced at 60° intervals in a hexagonal arrangement.

**Figure 4.**
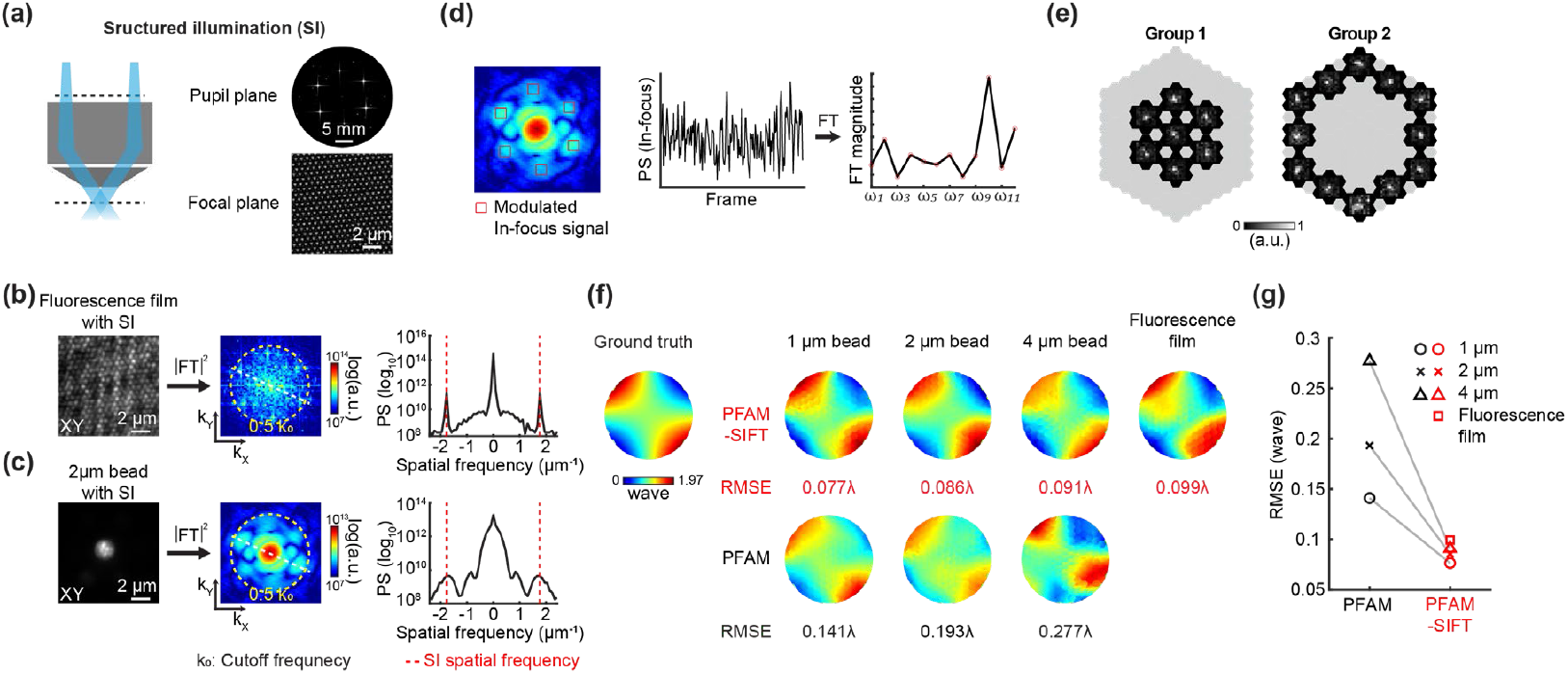
PFAM-SIFT enables accurate aberration measurement from spatially extended fluorescent structures. (**a**) (Left) Schematic of structured illumination (SI); (Right) Simulated illumination pattern at the back pupil plane of the microscope objective and the periodic hexagonal SI pattern at the objective focal plane. Yellow dashed circle: cutoff frequency; (**b**) (Left) Example image of a fluorescence film of 0.8 μm thickness under SI; (Middle) Power spectrum (PS) of the Fourier Transform (FT) of the image in logarithmic scale; (Right) Profile along the dashed white line in PS image. Yellow dashed circle: 0.5k_0_; k_0_: cutoff frequency; Red dashed lines: spatial frequency of SI. (**c**) Same as in (**b**) but for a 2-µm-diameter fluorescent bead. (**d**) PFAM-SIFT calculates the average PS values within regions containing modulated in-focus signals (red boxes) in each frame to obtain the PS trace, which is then Fourier transformed and Fourier magnitudes corresponding to the interference strength between each probe beamlet and the reference focus are extracted for phase gradient measurement. (**e**) Example tip/tilt interference maps obtained using PS values with two-group DM configuration. (**f**) (From left to right) Ground-truth (GT) corrective wavefronts for astigmatism applied to DM, corrective wavefronts from PFAM-SIFT (top) and PFAM (bottom) from 1 μm, 2 μm, and 4 μm fluorescent beads, and 0.8 μm fluorescence films, respectively, at a camera frame rate of 50 Hz. Each method was iterated three times. RMSE: RMS errors between GT and measured wavefronts. (**g**) RMSEs for PFAM and PFAM-SIFT from different fluorescent structures.

For a laterally extended structure such as a 0.8-μm-thick fluorescent film, SI generated fluorescence contrast within its XY image (left panel, **Fig. 4b**). As expected, the 2D Fourier power spectrum (i.e., Fourier magnitude squared) of the image had six peaks at the spatial frequency of SI (right panel and line profile, **Fig. 4b**). For a more axially extended structure such as a 2-μm-diameter fluorescent bead, in addition to generating contrast in the XY image (left panel, **Fig. 4c**), SI shifted in-focus signals into six side lobes centering at the SI spatial frequency in the Fourier domain (right panel and line profile, **Fig. 4c**); because out-of-focus fluorescence is largely unmodulated by SI, in the spatial frequency domain, it does not contribute to the power spectrum of these side lobes.

At each tip/tilt angles applied to a macro-segment during PFAM-SIFT, if the corresponding beamlet does not overlap the reference focus, modulating its phase would not change the Fourier power spectrum of these six side lobes. If the beamlet does overlap with the reference focus, phase modulation would lead to changes in the side-lobe power spectrum. By isolating and measuring how PFAM-SIFT modulates the total power spectrum within the side lobes, PFAM-SIFT achieves reliable aberration measurements from spatially extended fluorescent structures, even under strong background conditions (**Supplementary Note**).

From the camera frames recorded during PFAM-SIFT, we cropped out images centered on in-focus structures (typical image size: 128×128 pixels) and calculated their Fourier power spectrum. To isolate the modulated in-focus signal in the spatial frequency domain, we cropped six 7×7-pixel regions centered at the SI spatial frequency (red boxes in left panel, **Fig. 4d**) and computed the sum of their Fourier power spectrum. Instead of image pixel values as in PFAM, PFAM-SIFT then used the variations in this power spectrum sum (center panel, **Fig. 4d**) to extract the interference strengths for modulated beamlets (right panel, **Fig. 4d**) and acquire the tip/tilt map for each macro-segment using the same processing pipeline as in PFAM. Since PFAM-SIFT relies solely on the modulated signal, the effective signal strength is reduced compared to standard PFAM under the same illumination power. To enhance the modulated signal from the reference focus, we used a two-group DM configuration (**Fig. 4e**).

To validate the aberration measurement performance of PFAM-SIFT, we introduced artificial astigmatism (∼0.4 waves RMS) to the DM and compared the capability of PFAM and PFAM-SIFT in accurately measuring this aberration from images of 1 µm, 2 µm, and 4 µm beads, as well as a 0.8 µm-thick fluorescent thin film. Three iterations were performed for each method. We found that across all samples, wavefronts measured by PFAM-SIFT closely resembled the ground truth, with RMSE below 0.1 waves for all samples (upper panels, **Fig. 4f**). In contrast, corrective wavefronts from PFAM had increasingly large RMSEs with the increase in bead diameter, with wavefront measurement failed entirely for the fluorescent thin film (lower panels, **Fig. 4f**; **Fig. 4g**; **Supplementary Fig. 6**).

### Robust and parallel aberration measurement enables high-resolution imaging of mouse brain slices

We next applied PFAM-SIFT to measure aberrations from GFP-labeled neuronal processes and synapses within 100 µm-thick fixed mouse brain slices (Thy1-GFP line M). To introduce additional aberrations, the cover glass was tilted by 4° using a goniometer stage. Aberrations were measured from fluorescence signals in the region marked by the red box at a depth of 80 µm (“No AO”, **Fig. 5a**). To evaluate the accuracy of PFAM-SIFT in brain tissue, we compared the measured wavefronts with those obtained using DWS via a Shack–Hartmann sensor of a 2P guide star scanned over the same region. DWS was performed with one iteration, while PFAM-SIFT required two. The corrective wavefronts from both methods showed similar profile (**Fig. 5b**).

**Figure 5.**
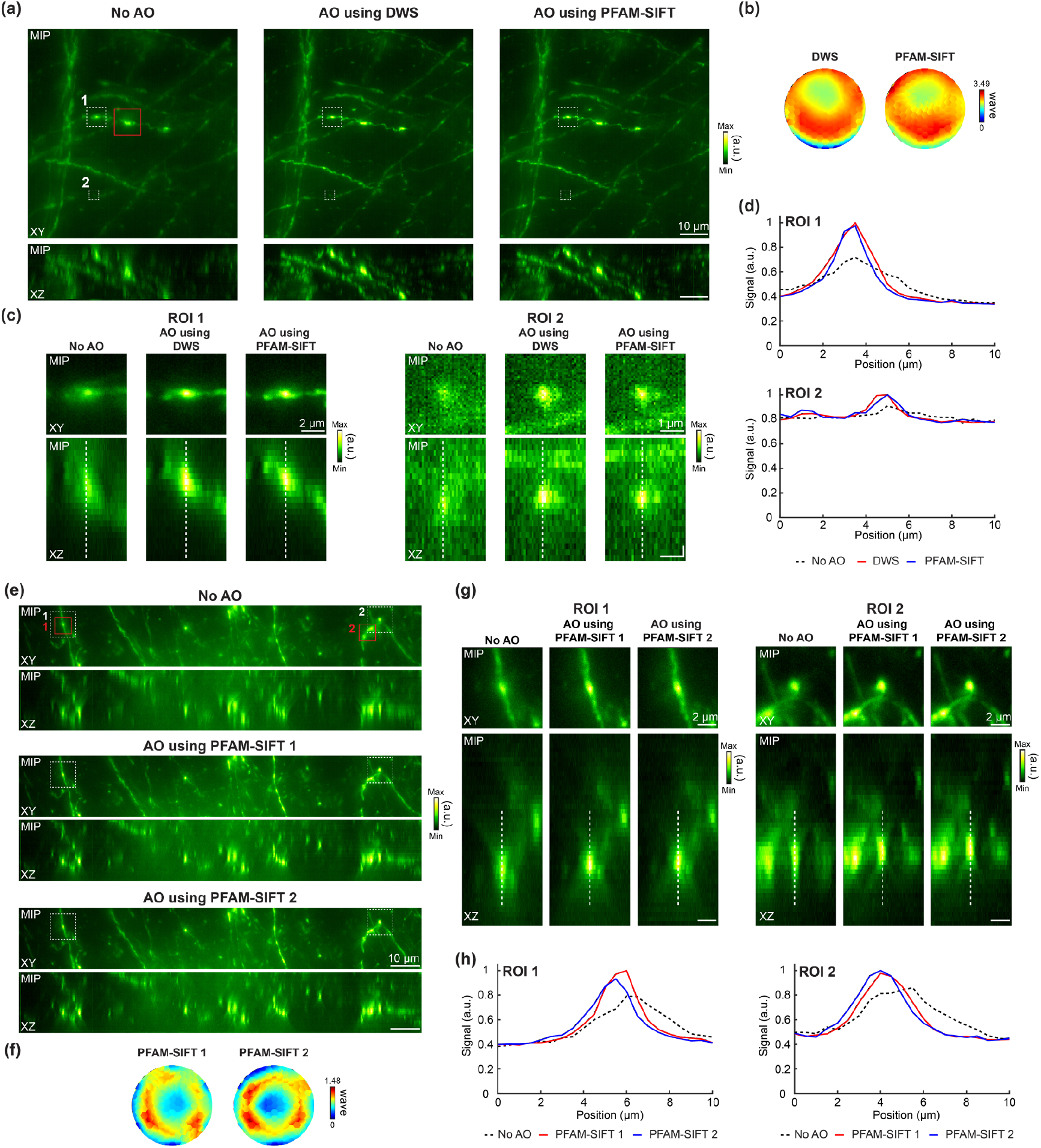
PFAM-SIFT improves widefield imaging quality of Thy1-GFP mouse brain slices. (**a**) XY and XZ maximum intensity projections (MIPs) from 76 μm × 76 μm × 20.5 μm widefield image stacks (80–100 μm imaging depth in slice; XY pixel size: 0.086 μm; Z pixel size: 0.5 μm) acquired without AO, with corrective wavefront from direct wavefront sensing (DWS) or from PFAM-SIFT. Both corrective wavefronts were measured from signals within the red box. DWS was iterated once, and PFAM-SIFT was iterated twice. All images are plotted with Greenhot color scale of the same minimal and maximal pixel values. (**b**) Corrective wavefronts from DWS and PFAM-SIFT. (**c**) Magnified XY and XZ MIPs of ROI 1 and 2 (white dashed boxes in (**a**)) acquired with No AO, with corrective wavefront from DWS, and with corrective wavefront from PFAM-SIFT. (**d**) Axial signal profiles along white dashed lines in (**c**). (**e**) PFAM-SIFT measures spatially varying aberrations in parallel from ROI 1 and 2 (red boxes). XY and XZ MIPs from 136 μm × 21 μm × 20.5 μm widefield image stacks (80–100 μm imaging depth; XY pixel size: 0.086 μm; Z pixel size: 0.5 μm) acquired with No AO, with corrective wavefront measured from ROI 1 (PFAM-SIFT 1), and with corrective wavefront measured from ROI 2 (PFAM-SIFT 2). PFAM-SIFT was iterated once. (**f**) Corrective wavefronts measured at ROI 1 and 2. (**g**) Magnified XY and XZ MIPs near ROI 1 and 2 (white boxes in (**e**)). (**h**) Axial signal profiles along white dashed lines in (**g**). PFAM-SIFT was performed at a camera frame rate of 50 Hz.

We then tested the performance of DWS and PFAM-SIFT by acquiring 20-µm-thick image stacks after applying their respective corrective wavefronts to the DM. In the resulting MIP images, both AO methods enhanced image resolution, contrast, and signal comparably over the entire FOV (**Fig. 5a**). These improvements enabled clearer visualization of fine neuronal structures such as axonal boutons (ROI 1, **Fig. 5c**) and dim dendritic spines (ROI 2, **Fig. 5c**), with their axial signal profiles having significantly increased contrast and peak fluorescence signal after correction (**Fig. 5d**). Fourier Ring Correlation (FRC) analysis^39^ further revealed comparable gains in spatial resolution (**Supplementary Fig. 7a**).

We next assessed the parallelized wavefront sensing capability of PFAM-SIFT in the same brain slice without tilting the cover glass. We measured aberrations from two ROIs (red boxes, “No AO”, **Fig. 5e**) and acquired their corresponding corrective wavefronts (**Fig. 5f**). These wavefronts were then separately applied to the DM for aberration correction, and the resulting widefield MIP images were compared. Both wavefronts led to global resolution improvement over the entire FOV (**Fig. 5e**). However, applying the corrective wavefront from ROI 1 (PFAM-SIFT 1) to the DM led to greater enhancement in ROI 1, while correction with ROI 2 wavefront (PFAM-SIFT 2) produced superior improvement in ROI 2. These region-specific enhancements were most evident in the XZ MIPs, where axial signal profiles became more symmetric (a sign for aberration-free imaging) and had stronger peak signals when the locally acquired corrective wavefront was applied (**Fig. 5g,h**). FRC analysis also confirmed that larger resolution gains were achieved with locally acquired corrective wavefronts (**Supplementary Fig. 7b**).

We were able to generate optically sectioned images through HiLo microscopy^40–42^, which computationally reconstructs an optically sectioned image from two widefield images acquired from the same structure: one acquired under uniform illumination and one under SI. A one-dimensional grating pattern was used for SI in HiLo microscopy. At each imaging depth, the optically sectioned images were computationally reconstructed using the standard HiLo reconstruction algorithm (Methods). Similar to our previous work showing that aberration correction improves optical-sectioning SIM images^14,15^, aberration correction using PFAM-SIFT improved HiLo image resolution and contrast as effectively as DWS (**Supplementary Fig. 8a-d**). Locally measured corrective wavefronts also led to better image quality (**Supplementary Fig. 8e-h**). Because PFAM-SIFT measurement uses in-focus signal specifically, which is given more weight in the HiLo reconstructed images, the enhancement of HiLo image quality was markedly more pronounced than for standard widefield images (cf., **Fig. 5** and **Supplementary Fig. 8**). The FOV-position-dependent improvement in image quality was also more pronounced in deconvolved images (**Supplementary Fig. 9**). Collectively, these results demonstrate that PFAM-SIFT enables accurate measurement and correction of spatially varying aberrations in complex biological tissues and facilitates parallel wavefront sensing across multiple locations from extended fluorescent features.

### PFAM-SIFT for in vivo structural imaging of the mouse brain

Finally, we applied PFAM-SIFT to in vivo structural imaging of neurons expressing tdTomato in the mouse brain. The mouse was anesthetized to minimize motion during aberration measurement. Due to photobleaching and sparse fluorescence structures within the FOV, PFAM-SIFT and DWS were performed with a single iteration at one FOV location (red box, “No AO”, **Fig. 6a**), yielding similar corrective wavefronts (**Fig. 6b**). The AO performance of each method was evaluated by acquiring 10 µm-thick image stacks without and with its corrective wavefront (**Fig. 6a**).

**Figure 6.**
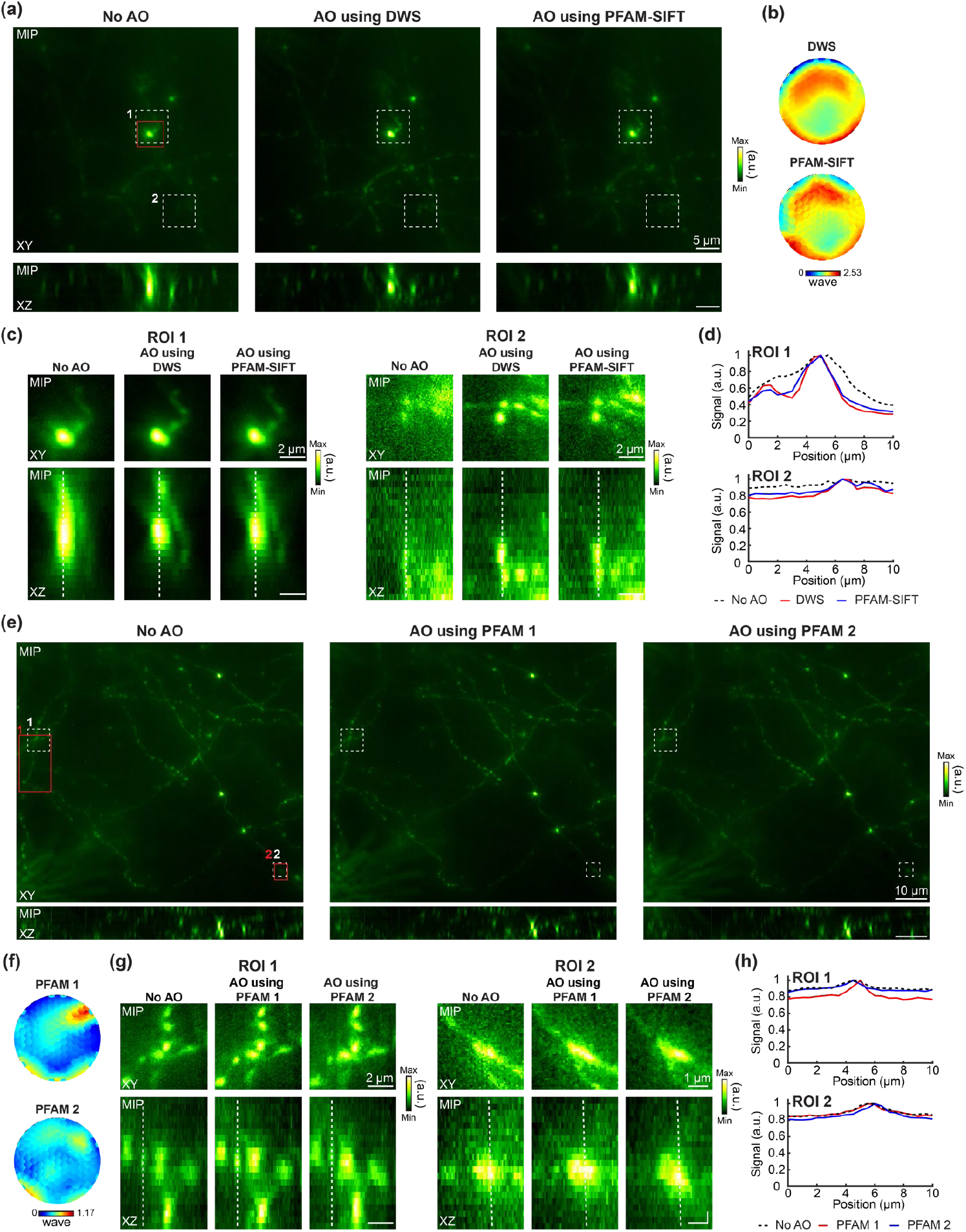
PFAM and PFAM-SIFT improve widefield imaging of tdTomato-expressing neurons in the mouse brain in vivo. (**a**) XY and XZ maximum-intensity projections (MIPs) from a 48 µm × 48 µm × 10.5 µm widefield image stack (25–35 µm below dura; XY pixel size: 0.086 μm; Z pixel size: 0.5 μm), acquired without AO, with corrective wavefront from direct wavefront sensing (DWS) or from PFAM-SIFT. Both corrective wavefronts were measured from signals within the red box. DWS and PFAM-SIFT were each iterated once. PFAM-SIFT was performed at a camera frame rate of 66 Hz. After registration in ImageJ (StackReg), images were individually normalized and plotted with Greenhot color scale. (**b**) Corrective wavefronts from DWS and PFAM-SIFT. (**c**) Magnified XY and XZ MIPs of ROIs 1 and 2 (white dashed boxes in (**a**)) acquired with No AO, with corrective wavefront from DWS, and with corrective wavefront from PFAM-SIFT. (**d**) Axial signal profiles along white dashed lines in (**c**), normalized by their maximum values. (**e**) PFAM of spatially varying aberrations in vivo from ROI 1 and 2 (red boxes) in parallel. XY and XZ MIPs from 85 µm × 94 µm × 10.5 µm widefield image stacks (25–35 µm below dura; XY pixel size: 0.086 μm; Z pixel size: 0.5 μm) acquired with No AO and with corrective wavefronts measured from ROI 1 (PFAM 1) or ROI 2 (PFAM 2) by PFAM at 0.172 µm pixel size using 2×2 camera pixel binning readout at a camera frame rate of 77 Hz. PFAM was iterated once. Images were registered and normalized as in (**a**). (**f**) Corrective wavefronts measured at ROIs 1 and 2. (**g**) Magnified XY and XZ MIPs near ROIs 1 and 2 (white boxes in (**e**)). (**h**) Axial signal profiles along white dashed lines in (**g**), normalized to their respective maxima.

Photobleaching reduced fluorescence signal intensity over time, making it difficult to compare image brightness across sequentially acquired stacks; therefore, individually normalized images were compared. Both PFAM-SIFT and DWS improved image resolution and contrast across the FOV (**Fig. 6a**) and in enlarged MIP views (**Fig. 6c**), with the resolution and contrast improvement especially striking along the axial direction as shown by the sharper axial signal profiles (**Fig. 6d**).

The dimmer fluorescence intensity and the sparser in vivo labeling led us to use PFAM instead of PFAM-SIFT for parallel aberration measurements, as PFAM-SIFT requires stronger fluorescence signals than found in the in vivo brain. With the FOV containing mostly sparse structures with dimensions comparable to the diffraction-limited focus, we used PFAM for parallel aberration measurements at two ROIs (red squares, “No AO”, **Fig. 6e**) across a large FOV. To accelerate acquisition over the large FOV, 2×2 camera pixel binning readout (effective pixel size: 0.176 μm) was applied for PFAM. Applying the corrective wavefronts (**Fig. 6f**) to the DM, we acquired widefield images (**Fig. 6e**). Although some residual aberrations remained—likely due to the use of PFAM instead of PFAM-SIFT—the application of the corrective wavefronts improved image contrast in the XZ MIP views for both ROIs (**Fig. 6g,h**). Notably, when the locally acquired corrective wavefront was applied to each ROI, the axial signal profiles became sharper (**Fig. 6h**).

We then applied patch-wise non-blind RLD to the ‘No AO’ image stack in **Fig. 6e** using the wavefront obtained at each location, allowing a direct comparison independent of photobleaching effects (**Supplementary Fig. 10**). The deconvolved images based on PFAM-SIFT showed slightly lower—but overall comparable—improvements in signal and contrast relative to those obtained using DWS. Furthermore, when region-specific corrective wavefronts from parallel PFAM were applied, the deconvolved images exhibited more pronounced improvements in axial signal contrast. These results demonstrate that both PFAM-SIFT and PFAM can successfully measure sample-induced aberrations in the mouse brain in vivo. Although here we only performed a single iteration due to photobleaching, we expect that additional iterations in more brightly labeled and photostable samples would yield further improvements in in vivo imaging quality.

## Discussion

In this study, we introduced PFAM, a multiplexed aberration measurement technique for widefield fluorescence microscopy. PFAM addresses two major limitations that have impeded the performance of AO in widefield imaging: the assumption of spatially invariant aberrations and the reliance on 3D-confined guide stars for accurate aberration measurement. By exploiting the highly parallel signal acquisition scheme of widefield detection cameras and fast segmented DMs that can be modulated at hundreds of Hz, PFAM enables massively parallel aberration measurement. It successfully measured complex, spatially varying aberrations at over 100 locations from the same FOV and enabled computational correction of image quality across the FOV via patch-wise non-blind RLD in a high-NA widefield microscope.

To improve aberration measurement accuracy using spatially extended fluorescent structures, we integrated PFAM with structured illumination (SI) and Fourier transform (FT) into a combined approach termed PFAM-SIFT. High-spatial-frequency SI generates contrast laterally in the focal plane and, for axially extended objects, shifts in-focus signals to the high-frequency region of the spatial frequency domain. By isolating and analyzing these modulated in-focus components, PFAM-SIFT enables accurate aberration measurement in extended samples, achieving similar performance to DWS without requiring additional optical sectioning techniques such as confocal^13^, two-photon^14,15^, or light-sheet excitation^43^. We validated PFAM-SIFT in both fixed and in vivo mouse brain tissues, demonstrating spatially varying aberration correction and image quality improvement across large FOVs.

While PFAM-SIFT broadens the applicability of PFAM, it requires relatively bright fluorescence signals. This limitation arises because PFAM-SIFT relies exclusively on the emitted fluorescence that is modulated by SI—i.e., the signal components shifted by SI (**Supplementary Note**). Under low signal-to-background ratio (SBR) conditions, these modulated signals may be too weak to support accurate wavefront measurements; In contrast, standard PFAM remains effective even under low-SBR conditions (**Supplementary Fig. 11**). Consequently, successful implementation of PFAM-SIFT requires bright fluorescence emission to ensure adequate signal strength.

Under current PFAM settings, total fluorescence acquisition time typically spans several minutes (Methods). However, this duration can be significantly reduced if fluorescence brightness or excitation power tolerable by the sample allows for shorter camera exposure times. For instance, with all other parameters held constant, reducing the exposure time to 1 ms—the minimum supported by current sCMOS cameras—would decrease the total acquisition time to 48.6 seconds for a 3-group DM configuration. This can be further reduced to 24.3 seconds or 12.1 seconds by lowering the sampling resolution to 2-group or 1-group DM configurations, respectively. In our current implementation, we group 7 segments of the 169-segment DM to form each macro-segment, resulting in a total of 18– 19 macro-segments. This suggests that PFAM can also be implemented by DMs with fewer segments (e.g., 37-segment DM from Boston Micromachines).

Currently, at each instant, only a single corrective wavefront can be physically applied to the DM, which optimally corrects aberration for a limited FOV. To physically correct spatially varying aberrations measured from multiple locations, methods such as sequentially switching between multiple corrective wavefronts^44^, multi-conjugate adaptive optics (MCAO)^45^, or multi-pupil adaptive optics (MPAO)^46^ can be employed. However, these approaches can increase imaging time, light exposure, or system complexity; none of them can be practically used to correct distinct aberrations at over >100 locations. Although PFAM and PFAM-SIFT can measure spatially varying aberrations simultaneously, optimal aberration correction over the whole FOV in this study were performed computationally using patch-wise non-blind deconvolution.

We anticipate that PFAM or PFAM-SIFT can be seamlessly integrated into a range of widefield-based imaging modalities, including SIM^13–15,18,37,47^, light-sheet fluorescence microscopy (LSFM)^43,48–50^, and single-molecule localization microscopy (SMLM)^17,51,52^. Given that SIM already incorporates structured illumination hardware, the implementation of PFAM-SIFT is both straightforward and synergistic. While we demonstrated the integration of PFAM-SIFT with optical-sectioning SIM, it can be readily extended to super-resolution SIM as well, for which an aberration-free imaging condition is even more important than for diffraction-limited imaging modalities^13,53^. LSFM provides intrinsic optical sectioning by illuminating a thin section using a light sheet, which effectively suppresses out-of-focus background. This makes LSFM highly compatible with standard PFAM, enabling large-FOV aberration correction without requiring PFAM-SIFT. In SMLM, accurate PSF characterization is essential for precise molecular localization. Spatially varying wavefronts measured by PFAM can be used to generate ROI-specific PSFs^54–56^, improving localization accuracy and extending the effective FOV. By combining PFAM with these advanced imaging platforms, AO can be more broadly deployed to support emerging biological applications that demand high resolution over large imaging areas.

In conclusion, PFAM and PFAM-SIFT offer practical and scalable solutions for correcting spatially varying aberrations in widefield fluorescence microscopy. By enabling robust, high-throughput, field-specific aberration measurement for diverse samples, they extend the reach of AO to widefield modalities where simultaneous multi-region correction is critical, advancing quantitative and high-resolution imaging of complex biological systems.

## Author Contribution

N.J. conceived of and supervised the project. H.K. designed and performed experiments; H.K. developed the PFAM algorithms; H.K. and I.K. analyzed the imaging data; R.N. prepared mouse samples; H.K., R.N., and N.J. wrote the manuscript with input from all authors.

## Acknowledgement

We thank Michael Helmbrecht for technical assistance and Qinrong Zhang for valuable advice on the optical system. This work was supported by NIH BRAIN® Initiative U01NS118300 and U01NS137449 (N.J.) and Weill Neurohub (N.J.).

## METHODS

### Nonbiological samples

Fluorescent beads with diameters of 0.2 µm, 0.5 µm, 1 µm, 2 µm (FluoSpheres™ Carboxylate-Modified Microspheres, yellow-green 505/515; ThermoFisher Scientific) or 4 µm (ThermoFisher Scientific TetraSpeck Microspheres, fluorescent blue/green/orange/dark red) were diluted 1:1000 to 1:3000 in deionized water and pipetted onto a microscope glass slide (12-550-12, Fisher Scientific). The slide was pre-coated with 10 mg/mL poly-L-lysine hydrobromide (P7890; Sigma-Aldrich) to help bead adhesion.

To prepare a thin fluorescent film, a highlighter pen was used to mark the surface of a glass slide. The thickness of the resulting fluorescent layer was measured by acquiring a z-stack using two-photon microscopy and using 0.5-µm-diameter beads as a reference.

To introduce spatially varying, sample-induced aberrations, a thin layer of transparent nail polish was applied to a No. 1 cover glass (Corning). This coated cover glass was then placed over the sample containing 0.5-µm-diameter fluorescent beads.

### Animals and surgical procedures

All animal experiments were conducted according to the National Institutes of Health guidelines for animal research. Procedures and protocols on mice were approved by the Institutional Animal Care and Use Committee at the University of California, Berkeley. Mice were housed in an animal facility at UC Berkeley campus with a 12-h light– dark cycle, ambient temperature between 20 and 26 °C, and humidity between 40 and 60%.

Brain slices were prepared from a Thy1-GFP line M transgenic mouse (Jackson Laboratory, stock no. 007788). Following deep anesthesia with isoflurane (Piramal), the animal underwent transcardial perfusion using phosphate-buffered saline (PBS; Invitrogen), followed by fixation with 4% paraformaldehyde (PFA; Electron Microscopy Sciences). The brain was then extracted and post-fixed overnight at 4 °C in a solution of 2% PFA and 15% sucrose in PBS. The next day, the solution was replaced with 30% sucrose in PBS, and the brain was stored at 4 °C for an additional 24 hours. Brain sections were obtained at a thickness of 100 µm using a microtome (Thermo Scientific Microm HM430). Slices were transferred into PBS, mounted onto microscope glass slides, and air-dried for approximately 2 hours. A No. 1.5 cover glass (Fisherbrand, 0.16–0.19 mm thickness) was applied using Vectashield Hardset Antifade mounting medium (H-1400). Imaging was performed once the mounting medium had fully cured.

In vivo imaging experiments were carried out on Thy1-GCaMP (GP5.17) mice (2–4 months old) sparsely expressing tdTomato. Cranial window implantation surgeries were performed under isoflurane anesthesia (1–2% in O_2_) following established procedures. Sparse expression of tdTomato was achieved in the mouse V1 L2/3 by injection of a 1:1 mixture of AAV2/1-CAG-FLEX-tdTomato (6.0 × 10^13^ GC per ml), and AAV2/1-hSyn-Cre (Addgene catalog #105553, diluted 10,000-fold to 3.2 × 10^8^ GC per ml by phosphate-buffered saline) in the Thy1-GCaMP mouse. 20– 50 nl of the virus mixture was injected at multiple sites 150–200 µm below the pia. A cranial window made of a glass coverslip (Fisher Scientific, no. 1.5) was embedded in the craniotomy and sealed in place with Vetbond tissue adhesive (3M). A metal headpost was then attached to the skull with Metabond, cyanoacrylate glue, and dental acrylic. In vivo imaging was carried out in the red fluorescence channel after 4 weeks of expression under isoflurane anesthesia (1% in O_2_).

### PFAM optical layout

The optical setup consisted of two optical paths (**Supplementary Fig. 2**): one for widefield fluorescence microscopy and the other for the TPE path used for DWS. A 488-nm or 561-nm continuous-wave laser was reflected off a spatial light modulator (SLM; Forth Dimension Displays Ltd., QXGA-3DM), which was optically conjugated to the objective focal plane. An achromatic half-wave plate (HWP; Bolder Vision Optik, AHWP3), placed between the SLM and a polarizing beam splitter (PBS; Thorlabs, PBS251), rotated the beam’s polarization to maximize the diffraction efficiency of the SLM. The SLM operated in two modes: SI and uniform illumination. In SI mode, a high-frequency hexagonal lattice or one-dimensional grating pattern displayed on the SLM generated a first-order diffracted beam, which was focused onto the front focal plane of a lens (L1; FL = 150 mm). This beam was relayed to the back focal plane of a water-immersion objective lens (Nikon, CFI Apo LWD 25×, NA 1.1, WD 2 mm), and interference between diffracted beams formed a hexagonal lattice or sinusoidal illumination pattern at the sample plane. In uniform illumination mode, the SLM was set to a flat phase pattern, acting as a mirror, and the beam followed the same optical path to the objective lens. Fluorescence emitted by the sample was collected by the objective and transmitted through a dichroic mirror (D1; Semrock, Di-R405/488/561/635-t3-25×36), which reflected the illumination light. The back focal plane of the objective was conjugated to a deformable mirror (DM) via a relay lens system (L4–L5; FL = 400 mm and 175 mm). The fluorescence reflected by the DM was then focused onto a widefield camera (Hamamatsu, ORCA Flash 4.0) through a series of lenses (L6, L7, L8; FL = 300 mm, 85 mm, and 75 mm, respectively).

For the TPE path used for DWS, a two-photon excitation beam was used to generate a two-photon-excited guide star. The output beam from a Ti:sapphire laser (Coherent, Chameleon Ultra II) was tuned to 920 nm or 1000 nm and scanned using a pair of galvanometric mirrors, which were conjugated via an achromatic lens pair (L11–L12; FL = 85 mm). This scanned beam was further relayed to the DM through another lens pair (L9–L10; FL = 300 mm and 85 mm). A movable mirror (MM) was inserted into the TPE path to redirect the two-photon excitation beam into the widefield system. This MM was mounted on a nanopositioning stage (SmarAct, SLC-24105-D-SC) for precise translation. The two-photon-excited fluorescence signal followed the same return path as the widefield emission. After being reflected by the DM and MM and descanned by the galvanometers, the beam was directed by a dichroic mirror (D2; Semrock, Di03-R785-t3-25×36) toward a Shack–Hartmann (SH) sensor. The SH sensor consisted of a lenslet array (Advanced Microoptic Systems GmbH) and a camera (Hamamatsu, ORCA Flash 4.0) positioned at the focal plane of the lenslet array.

System aberrations were corrected prior to sample-induced aberration measurements. Additionally, the objective correction collar was adjusted to compensate for spherical aberrations introduced by the cover glass and cranial window during brain tissue and in vivo imaging. These conditions are referred to as “No AO.” The measured post-objective laser power varied depending on the sample: 48 μW to 0.3 mW for fluorescent beads, 72.6 μW to 135 μW for brain slices, and 100 μW for in vivo brain samples.

### Implementation and parameters of PFAM

Our DM segmented the pupil into 169 segments. To enhance modulation strength, we grouped 7 adjacent segments and controlled their tip, tilt and piston values so that they formed a macro-segment with a continuous surface.

During PFAM, the DM was divided into macro-segments that were modulated (gold segments in Step 1, **Supplementary Fig. 1**) and segments that were fixed (gray segments in Step 1, **Supplementary Fig. 1**). Light reflected off fixed segments formed a reference focus on the camera. Then, specific tip angles Θ_*i*_ and tilt angles Φ_*j*_ (*i, j* = 1, 2, …, *n*) were applied to the macro-segments (gold segments in **Supplementary Fig. 1**), with each angle randomly selected from a list of *n* angles evenly spaced between –Ψ/2 and Ψ/2. The applied tip and tilt angles displaced the focus of the corresponding probe beamlet along the X and Y axes in the image plane by X_*i*_ and Y_*j*_, respectively, with X_*i*_ = *f* × tan(2Θ_*i*_/M), Y_*j*_ = *f* × tan(2Φ_*j*_ /M). Here, *f* is the focal length of the tube lens (in our system, L8), and M is the magnification from the DM to the pupil plane of the tube lens. We then simultaneously modulated the piston value of each macro-segment at a distinct frequency ω_*s*_ and recorded the fluorescence images over N_*f*_ frames (left panel, Step 2, **Supplementary Fig. 1**). We then averaged the pixel values from a target image area (e.g., red box, left panel, Step 2, **Supplementary Fig. 1**) and plotted the average values over time (right panel, Step 2, **Supplementary Fig. 1**). This trace was then Fourier transformed (Step 3, **Supplementary Fig. 1**), and the Fourier magnitude at the modulation frequency ω_*s*_ (red circles, Step 3, **Supplementary Fig. 1**) indicated the interference strength between the corresponding probe beamlet at the given displacement (X_*i*_, Y_*j*_) and the reference focus.

This procedure was repeated *n* × *n* times to sample all tip/tilt angle combinations within [–Ψ/2, Ψ/2], effectively scanning every probe focus around the reference focus over 2*f* × tan(Ψ/M) along both X and Y (Step 4, **Supplementary Fig. 1**). For each segment, we calculated the Fourier magnitude versus displacement (X_*i*_, Y_*j*_) in the camera plane, producing a tip/tilt interference map, which revealed how interference strength depends on probe beamlet displacement. For visualization, the tip/tilt interference map was scaled to the sample plane coordinates (Step 5, **Supplementary Fig. 1**). The centroid of the tip/tilt map corresponds to the tip/tilt that needed to be added to each modulated macro-segment in order for the foci of their corresponding probe beamlets to maximally interfere with the reference focus.

When needed, we repeated steps 1–5 for different macro-segment groups (Step 6, **Supplementary Fig. 1**). The total fluorescence signal acquisition time was *m* × *n* × *n* × N_*f*_ × *t*, where *m* is the number of groups and *t* is the exposure time per frame.

A representative set of parameters used for PFAM with fluorescent beads is as follows: 200 frames (N_*f*_ = 200) were captured for each of the 9 × 9 grid of tip and tilt angles (Ψ = 1.2 mrad, *n* = 9), with a camera exposure time of 8 ms. For brain slice and in vivo brain imaging, the camera exposure time was increased to 20 ms and 15 ms, respectively, due to lower fluorescence brightness.

In the 3-groups DM configuration used for bead measurements, the total image acquisition time was 388.8 s, whereas the 2-groups and 1-group DM configurations required 259.2 s and 129.6 s, respectively (**Supplementary Fig. 3**). For brain slice and in vivo experiments, the 2-group configuration was used, resulting in a total acquisition time of 648 s and 486 s, respectively.

Photobleaching during aberration measurement was approximately 1–10% for brain slices and 5–7% for in vivo imaging. During tip/tilt scanning, the tip and tilt angles were applied in random order. As a result, the reduction in signal due to photobleaching increased noise in the tip/tilt interference map but did not bias the centroid location in the map.

### Operation parameters for PFAM-SIFT

In PFAM-SIFT, the sample was illuminated with hexagonal lattice pattern. The spatial frequency of the stripes was set to 0.43 k_0_ for 0.5-µm, 1-µm, 2-µm-diameter bead samples and 0.29 k_0_ for 4-µm-diameter bead samples, brain slice and in vivo brain imaging, respectively, where k_0_ is the cutoff frequency of the optical system. Here, due to the strong out-of-focus signal relative to the in-focus signal in 4-µm-diameter beads and biological tissues, a lower spatial frequency was used to increase the depth of field of the modulation, thereby enhancing the detectability of the modulated signal over background fluorescence.

### HiLo reconstruction

HiLo reconstruction was performed using an ImageJ plugin ‘HiLo’ developed by the laboratory of Jerome Mertz (http://biomicroscopy.bu.edu/). The SI pattern was a sinusoidal grating with a spatial frequency of 0.58 k_0_ at the sample plane, where k_0_ is the cutoff spatial frequency of the optical system.

The HiLo plugin requires several input parameters: the scaling factor, camera gain, and readout noise for shot noise bias correction. The HiLo scaling factor, which scales the low spatial frequency component to balance with the high-frequency component, was set to 1.1. For shot noise bias correction, we used the camera specifications of the Hamamatsu ORCA-Flash4.0 (C11440-22CU), with a gain of 0.46 electrons/ADU and a readout noise of 1.6 electrons (rms).

### Direct wavefront sensing

For brain slice and in vivo mouse brain samples, ground-truth sample-induced aberrations were acquired using DWS. Prior to measuring sample-induced aberrations, system aberrations in the TPE path were characterized and corrected via modal AO^31^. Individual Zernike modes were sequentially applied to the DM to determine the corrective wavefront for system aberration. After applying the system-correction wavefront to the DM, the TPE beam was scanned over a 4.8 µm × 4.8 µm region containing a 2-µm-diameter fluorescent bead and the pattern on the SH sensor was recorded as the aberration-free reference. To measure sample-induced aberrations, a 10 µm × 10 µm region centered on an isolated fluorescent structure was selected, and a new SH pattern was acquired. Spot displacements in the new SH pattern relative to the reference pattern were converted into local wavefront slopes and used to computationally reconstruct the continuous aberrated wavefront, assuming a continuous phase profile. The wavefront reconstruction method used in DWS was the same as that used for PFAM.

### Phase retrieval

In widefield microscopy, ground truth system aberrations were determined by phase retrieval with the Gerchberg– Saxton algorithm^36^ applied to a 3‐D stack of 0.2-µm-diameter fluorescent beads. The phase retrieval was performed with two iterations for system aberrations (**Fig. 1**).

### Non-blind Richardson-Lucy deconvolution

Non-blind deconvolution was performed using a GPU-accelerated Python implementation of the Richardson–Lucy algorithm (RLD)^57^. The deconvolution was applied to 3-D stack images acquired without AO correction, using 3D PSFs generated from the measured wavefront aberrations. A machine with a NVIDIA A40 and an Intel Xeon Gold 6354 CPU was used for deconvolution. For bead samples, 50 iterations (7 s of computation for **Fig. 3, Supplementary Fig. 4**) were used; 100 iterations (13 s of computation for **Supplementary Fig. 9**) were applied to brain slice samples, and 150 iterations (12 s of computation for **Supplementary Fig. 10**) to in vivo brain samples. In patch-wise non-blind RLD, fluorescent beads were first clustered based on spatial proximity, and the entire FOV was divided into multiple patches. Clustering was performed using the clusterdata() function from the MATLAB Statistics and Machine Learning Toolbox. Each patch was then deconvolved using the wavefront computed from the average signal variations of all beads within that patch.

## References

1. Ji, N. Adaptive optical fluorescence microscopy. Nat Methods 14, 374–380 (2017).

2. Hampson, K. M. et al. Adaptive optics for high-resolution imaging. Nature Reviews Methods Primers 1, 68 (2021).

3. Zhang, Q. et al. Adaptive optics for optical microscopy. Biomed Opt Express 14, 1732–1756 (2023).

4. Kam, Z., Hanser, B., Gustafsson, M. G. L., Agard, D. A. & Sedat, J. W. Computational adaptive optics for live three-dimensional biological imaging. Proceedings of the National Academy of Sciences 98, 3790–3795 (2001).

5. Hom, E. F. Y. et al. AIDA: an adaptive image deconvolution algorithm with application to multi-frame and three-dimensional data. Journal of the Optical Society of America A 24, 1580–1600 (2007).

6. Kohli, A. et al. Ring deconvolution microscopy: exploiting symmetry for efficient spatially varying aberration correction. Nat Methods 1–10 (2025).

7. Hu, Q. et al. Universal adaptive optics for microscopy through embedded neural network control. Light Sci Appl 12, 270 (2023).

8. Kang, I., Zhang, Q., Yu, S. X. & Ji, N. Coordinate-based neural representations for computational adaptive optics in widefield microscopy. Nat Mach Intell 6, 714–725 (2024).

9. Guo, M. et al. Deep learning-based aberration compensation improves contrast and resolution in fluorescence microscopy. Nat Commun 16, 313 (2025).

10. Platt, B. C. & Shack, R. History and principles of Shack-Hartmann wavefront sensing. Journal of refractive surgery vol. 17 S573–S577 Preprint at (2001).

11. Azucena, O. et al. Adaptive optics wide-field microscopy using direct wavefront sensing. Opt Lett 36, 825–827 (2011).

12. Jorand, R. et al. Deep and clear optical imaging of thick inhomogeneous samples. PLoS One 7, e35795 (2012).

13. Lin, R., Kipreos, E. T., Zhu, J., Khang, C. H. & Kner, P. Subcellular three-dimensional imaging deep through multicellular thick samples by structured illumination microscopy and adaptive optics. Nat Commun 12, 3148 (2021).

14. Zhang, Q., Pan, D. & Ji, N. High-resolution in vivo optical-sectioning widefield microendoscopy. Optica 7, 1287–1290 (2020).

15. Li, Z. et al. Fast widefield imaging of neuronal structure and function with optical sectioning in vivo. Sci Adv 6, eaaz3870 (2020).

16. Scrimgeour, J. & Curtis, J. E. Aberration correction in wide-field fluorescence microscopy by segmented-pupil image interferometry. Opt Express 20, 14534–14541 (2012).

17. Burke, D., Patton, B., Huang, F., Bewersdorf, J. & Booth, M. J. Adaptive optics correction of specimen-induced aberrations in single-molecule switching microscopy. Optica 2, 177–185 (2015).

18. Žurauskas, M. et al. IsoSense: frequency enhanced sensorless adaptive optics through structured illumination. Optica 6, 370–379 (2019).

19. Hanser, B. M., Gustafsson, M. G. L., Agard, D. A. & Sedat, J. W. Phase retrieval for high-numerical-aperture optical systems. Opt Lett 28, 801–803 (2003).

20. Hanser, B. M., Gustafsson, M. G. L., Agard, D. A. & Sedat, J. W. Phase‐retrieved pupil functions in wide‐ field fluorescence microscopy. J Microsc 216, 32–48 (2004).

21. Kner, P., Winoto, L., Agard, D. A. & Sedat, J. W. Closed loop adaptive optics for microscopy without a wavefront sensor. in Proceedings of SPIE vol. 7570 10–1117 (2010).

22. Park, J.-H., Sun, W. & Cui, M. High-resolution in vivo imaging of mouse brain through the intact skull. Proceedings of the National Academy of Sciences 112, 9236–9241 (2015).

23. Li, J. et al. Conjugate adaptive optics in widefield microscopy with an extended-source wavefront sensor. Optica 2, 682–688 (2015).

24. Mertz, J., Paudel, H. & Bifano, T. G. Field of view advantage of conjugate adaptive optics in microscopy applications. Appl Opt 54, 3498–3506 (2015).

25. Ancora, D., Furieri, T., Bonora, S. & Bassi, A. Spinning pupil aberration measurement for anisoplanatic deconvolution. Opt Lett 46, 2884–2887 (2021).

26. Furieri, T. et al. Aberration measurement and correction on a large field of view in fluorescence microscopy. Biomed Opt Express 13, 262–273 (2021).

27. Wu, T., Guillon, M., Tessier, G. & Berto, P. Multiplexed wavefront sensing with a thin diffuser. Optica 11, 297–304 (2024).

28. Richardson, W. H. Bayesian-based iterative method of image restoration. J Opt Soc Am 62, 55–59 (1972).

29. Lucy, L. B. An iterative technique for the rectification of observed distributions. Astronomical Journal, Vol. 79, p. 745 (1974) 79, 745 (1974).

30. Panagopoulou, S. I. & Neal, D. P. Zonal matrix iterative method for wavefront reconstruction from gradient measurements. Journal of Refractive Surgery vol. 21 S563–S569 Preprint at (2005).

31. Zhang, Q. et al. Retinal microvascular and neuronal pathologies probed in vivo by adaptive optical two-photon fluorescence microscopy. Elife 12, e84853 (2023).

32. Wang, C. et al. Multiplexed aberration measurement for deep tissue imaging in vivo. Nat Methods 11, 1037–1040 (2014).

33. Rodríguez, C. et al. An adaptive optics module for deep tissue multiphoton imaging in vivo. Nat Methods 18, 1259–1264 (2021).

34. Rodríguez, C. et al. Adaptive optical third-harmonic generation microscopy for in vivo imaging of tissues. Biomed Opt Express 15, 4513 (2024).

35. Pan, D. et al. Frequency-multiplexed aberration measurement for confocal microscopy. Opt Express 32, 28655–28665 (2024).

36. Gerchberg, R. W. A practical algorithm for the determination of plane from image and diffraction pictures. Optik (Stuttg) 35, 237–246 (1972).

37. Débarre, D., Botcherby, E. J., Booth, M. J. & Wilson, T. Adaptive optics for structured illumination microscopy. Opt Express 16, 9290–9305 (2008).

38. Pedrazzani, M. et al. Sensorless adaptive optics implementation in widefield optical sectioning microscopy inside in vivo Drosophila brain. J Biomed Opt 21, 36006 (2016).

39. Descloux, A., Grußmayer, K. S. & Radenovic, A. Parameter-free image resolution estimation based on decorrelation analysis. Nat Methods 16, 918–924 (2019).

40. Lim, D., Chu, K. K. & Mertz, J. Wide-field fluorescence sectioning with hybrid speckle and uniform-illumination microscopy. Opt Lett 33, 1819–1821 (2008).

41. Mertz, J. & Kim, J. Scanning light-sheet microscopy in the whole mouse brain with HiLo background rejection. J Biomed Opt 15, 16027 (2010).

42. Lim, D., Ford, T. N., Chu, K. K. & Mertz, J. Optically sectioned in vivo imaging with speckle illumination HiLo microscopy. J Biomed Opt 16, 16014 (2011).

43. Hubert, A. et al. Adaptive optics light-sheet microscopy based on direct wavefront sensing without any guide star. Opt Lett 44, 2514–2517 (2019).

44. Wang, K. et al. Rapid adaptive optical recovery of optimal resolution over large volumes. Nat Methods 11, 625–628 (2014).

45. Simmonds, R. D. & Booth, M. J. Modelling of multi-conjugate adaptive optics for spatially variant aberrations in microscopy. Journal of Optics 15, 094010 (2013).

46. Park, J.-H., Kong, L., Zhou, Y. & Cui, M. Large-field-of-view imaging by multi-pupil adaptive optics. Nat Methods 14, 581–583 (2017).

47. Thomas, B., Wolstenholme, A., Chaudhari, S. N., Kipreos, E. T. & Kner, P. Enhanced resolution through thick tissue with structured illumination and adaptive optics. J Biomed Opt 20, 26006 (2015).

48. Bourgenot, C., Saunter, C. D., Taylor, J. M., Girkin, J. M. & Love, G. D. 3D adaptive optics in a light sheet microscope. Opt Express 20, 13252–13261 (2012).

49. Liu, T.-L. et al. Observing the cell in its native state: Imaging subcellular dynamics in multicellular organisms. Science (1979) 360, eaaq1392 (2018).

50. Mcfadden, C. et al. Adaptive optics in an oblique plane microscope. Biomed Opt Express 15, 4498–4512 (2024).

51. Xu, F. et al. Three-dimensional nanoscopy of whole cells and tissues with in situ point spread function retrieval. Nat Methods 17, 531–540 (2020).

52. Zhang, P. et al. Deep learning driven adaptive optics for single molecule localization microscopy. Biophys J 122, 430a (2023).

53. Turcotte, R. et al. Dynamic super-resolution structured illumination imaging in the living brain. Proceedings of the National Academy of Sciences 116, 9586–9591 (2019).

54. Xiao, D. et al. Large-FOV 3D localization microscopy by spatially variant point spread function generation. Sci Adv 10, eadj3656 (2024).

55. Liu, S. et al. Universal inverse modeling of point spread functions for SMLM localization and microscope characterization. Nat Methods 21, 1082–1093 (2024).

56. Droste, I. et al. Calibration-free estimation of field-dependent aberrations for single-molecule localization microscopy across large fields of view. Optica 12, 1220–1229 (2025).

57. Perdigão, L. M. A., Berger, C., Yee, N. B. Y., Darrow, M. C. & Basham, M. RedLionfish–fast Richardson-Lucy Deconvolution package for efficient point spread function suppression in volumetric data. Wellcome Open Res 9, 296 (2024).

