## Supplementary Material for "Parallel frequency-multiplexed aberration measurement for widefield fluorescence microscopy"

### Supplementary Note.

#### Theoretical background of PFAM-SIFT

In PFAM-SIFT, high-spatial-frequency illumination patterns predominantly modulate the in-focus signal, while the out-of-focus background fluorescence remains largely unmodulated. For simplicity, we consider a 1D sample illuminated with a sinusoidal pattern of spatial frequency $k_g$. The detected signal can be written as:

$$S(x) = \frac{1}{2}\left[I_{\text{in}}(x)\left(1 + M\sin\left(k_g x\right)\right) + I_{\text{out}}(x)\right],$$

where $I_{\text{in}}(x)$ and $I_{\text{out}}(x)$ represent the in-focus and out-of-focus signal components, respectively, and $M$ is the modulation depth. Taking the Fourier transform of $S(x)$, we obtain:

$$\tilde{S}(k) = \frac{1}{2}\left[\tilde{I}_{\text{in}}(k) + \tilde{I}_{\text{out}}(k) + \frac{M}{2i}\left(\tilde{I}_{\text{in}}\left(k - k_g\right) - \tilde{I}_{\text{in}}\left(k + k_g\right)\right)\right].$$

In this expression, only the in-focus component $\tilde{I}_{\text{in}}$ appears at the modulation sidebands $k \pm k_g$, while the out-of-focus component remains centered at low spatial frequencies due to its inherently low-frequency spatial structure. Therefore, by isolating the sideband signals $\tilde{I}_{\text{in}}\left(k \pm k_g\right)$, we can effectively enhance the in-focus signal over the out-of-focus background. Wavefront sensing is then performed based on the power spectrum which is the squared Fourier magnitude at these modulation frequencies.

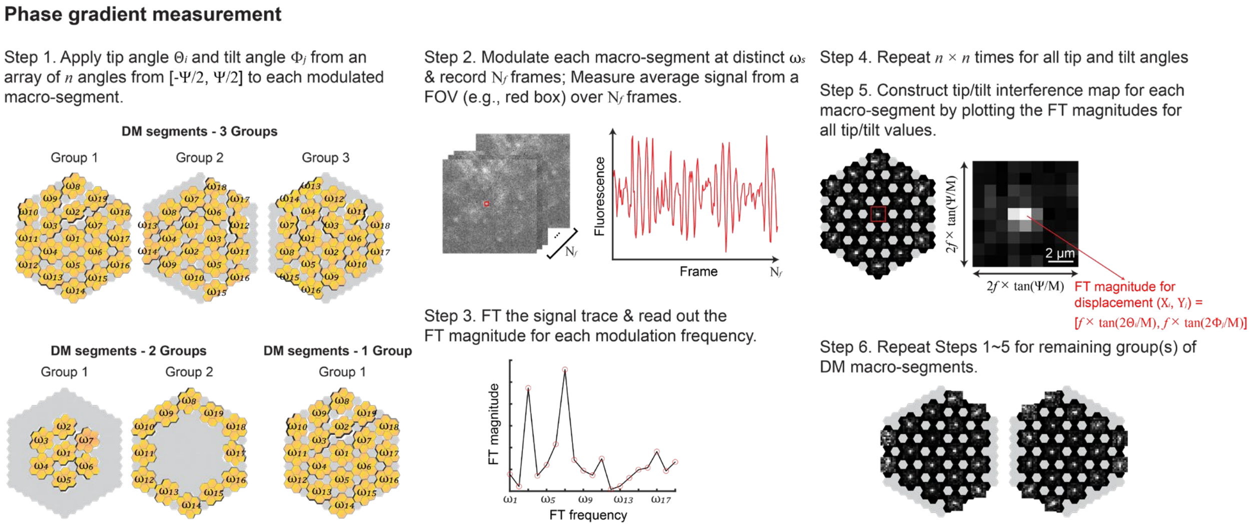


**Supplementary Figure 1. Schematics of phase gradient measurement in PFAM.** Step 1. Select DM configuration of 3 groups, 2 groups, or 1 group. Tip and tilt values are applied to modulated macro-segments (gold), while fixed segments (gray) remain static. Tip angles $\Theta_i$ and tilt angles $\Phi_j$ ($i$, $j$ = 1, 2, ..., $n$) are randomly selected from a list of $n$ angles evenly spaced between –Ψ/2 and Ψ/2. The piston value of each segment is modulated at a unique frequency $\omega_s$. Step 2. At each tip/tilt angle set, $N_f$ frames are recorded by the widefield camera during piston modulation, and average signal from a select region (e.g., red box) is measured across the recorded frames. Step 3. The recorded signal trace is Fourier transformed and the Fourier magnitudes at each $\omega_s$ are extracted, reflecting the interference strength between each probe beamlet and the reference focus under this set of tip/tilt values. Step 4. This process is repeated $n \times n$ times to cover all combinations of tip and tilt values. Step 5. For each modulated macro-segment, its Fourier magnitude is plotted as a function of focal displacements ($X_i$, $Y_j$), resulting in a 2D tip/tilt interference map that represents the interference strength between each probe beamlet and the reference focus (Methods). Here, interference maps are scaled to the object plane for presentation purposes. Step 6. Steps 1–5 are repeated for different combinations of fixed and modulated segment groups.

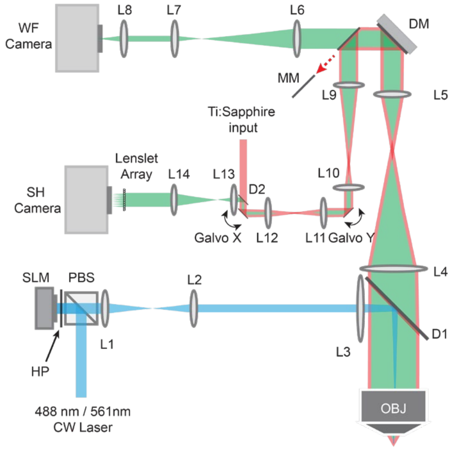


**Supplementary Figure 2. Optical setup for PFAM and DWS with two-photon excitation (TPE).** PBS: polarizing beam splitter; HP: half-wave plate; SLM: spatial light modulator; L, lenses; D1, D2: dichroic mirrors; OBJ, objective lens; DM: deformable mirror; MM, movable mirror; WF: widefield; Galvo X and Y, galvanometers; SH, Shack-Hartman; Blue path: excitation light for widefield illumination; Red path: excitation light for two-photon fluorescence guide star generation; Green path: fluorescence emission.

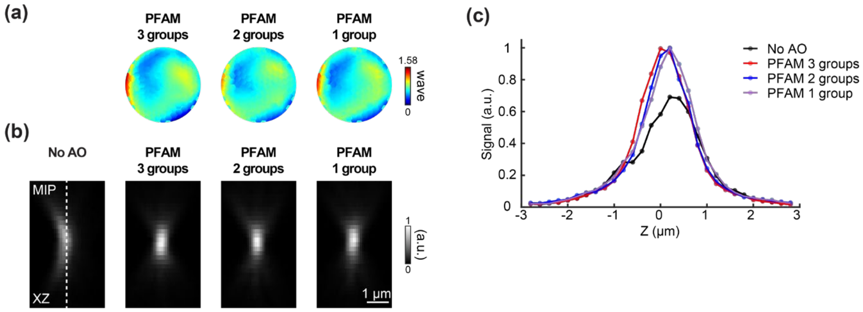


**Supplementary Figure 3. System aberration correction using PFAM with different DM grouping strategies.** System aberrations were measured with three different DM grouping configurations: 3 groups, 2 groups, and 1 group (Step 1, **Supplementary Fig. 1**). (**a**) Corrective wavefronts measured using each DM configuration after two iterations. (**b**) Maximum intensity projection (MIP) images in XZ of a 0.5-μm-diameter fluorescent bead acquired under each correction condition. (**c**) Axial intensity profiles along dashed lines in (**b**).

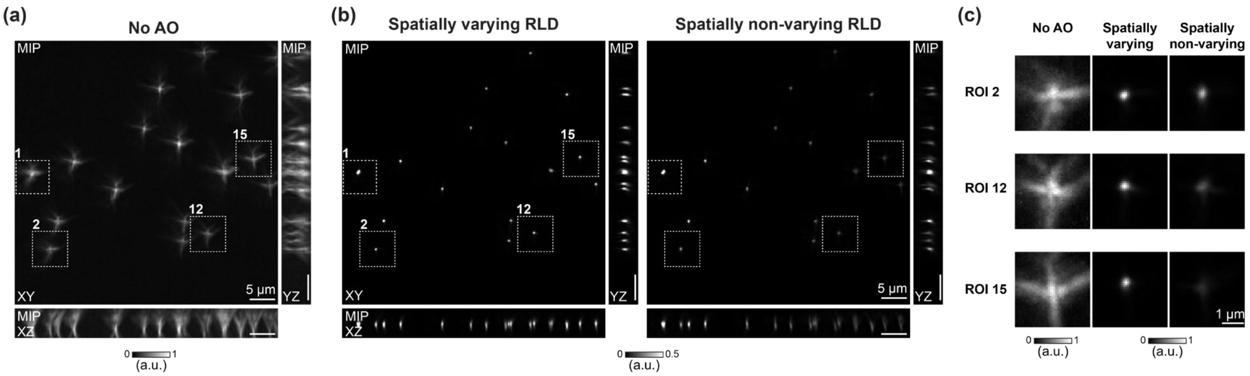


**Supplementary Figure 4. PFAM measurement and Richardson-Lucy deconvolution of spatially varying sample-induced aberrations.** Same data as **Fig. 3h-k**, after Richardson–Lucy deconvolution (RLD) using spatially varying or non-varying aberration. (**a**) XY, XZ, and YZ MIPs of 0.5-µm-diameter fluorescent beads with spatially varying aberrations. (**b**) XY, XZ, and YZ MIPs after RLD using spatially varying wavefronts ("Spatially varying RLD") or the corrective wavefront measured at ROI 1 ("spatially non-varying RLD"). Images are saturated for visibility. (**c**) Magnified XY MIPs at ROIs 2, 12, 15 of "No AO" images, deconvolved images with spatially varying wavefronts, and deconvolved images with wavefront measured at ROI 1. Gray scale for 'No AO' image is normalized to its minimal and maximal pixel values, while deconvolved images share the same minimal and maximal values for gray scale.

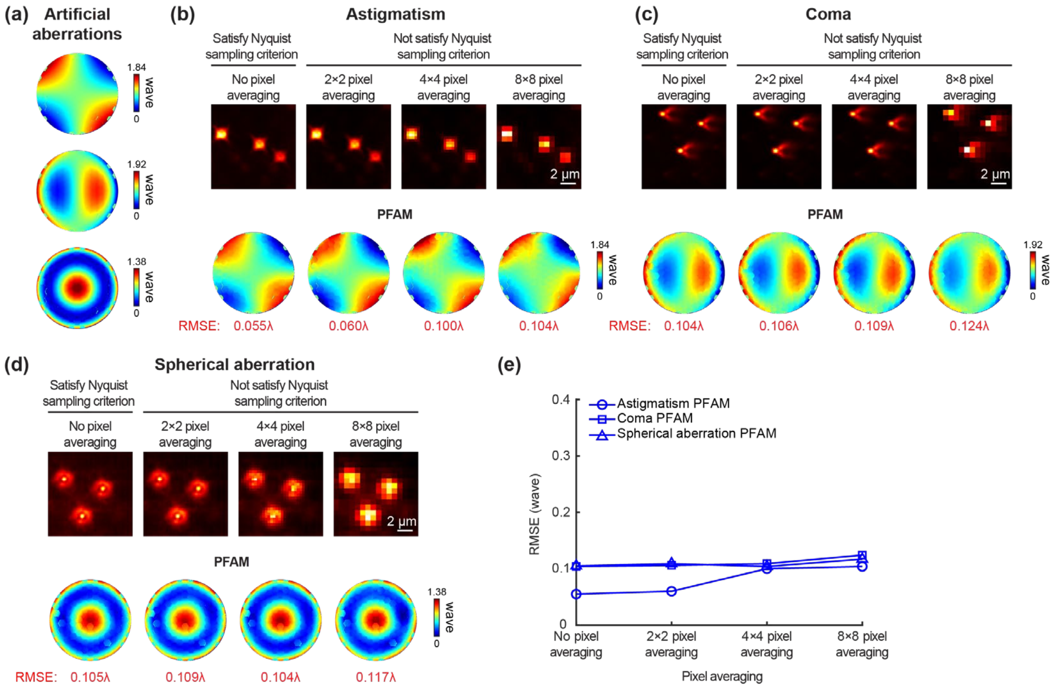


**Supplementary Figure 5. PFAM does not require camera pixel size to satisfy Nyquist sampling criterion.** (**a**) Ground-truth (GT) corrective wavefronts for artificial aberrations applied to DM: (top to bottom) astigmatism of 0.37 waves RMS, coma of 0.36 waves RMS, and spherical aberration of 0.36 waves RMS. (**b**) (Top) XY images of 0.5-μm-diameter beads acquired with applied astigmatism with pixel sizes of 0.086 μm, 0.172 μm, 0.344 μm, and 0.688 μm, respectively, with larger pixel sizes achieved by 2×2, 4×4, and 8×8 pixel averaging on the image of 0.086 μm pixel size; Nyquist sampling pixel size: 0.117 μm (for 0.234 μm lateral resolution). (Bottom) Corrective wavefronts measured by 1-iteration of PFAM from images with pixel sizes above; RMSE: RMS errors between GT and measured wavefronts. (**c**) Same as (**a**), but for coma. (**d**) Same as (**a**), but for spherical aberration. (**e**) RMSE between GT wavefronts and measured wavefronts across pixel averaging conditions.

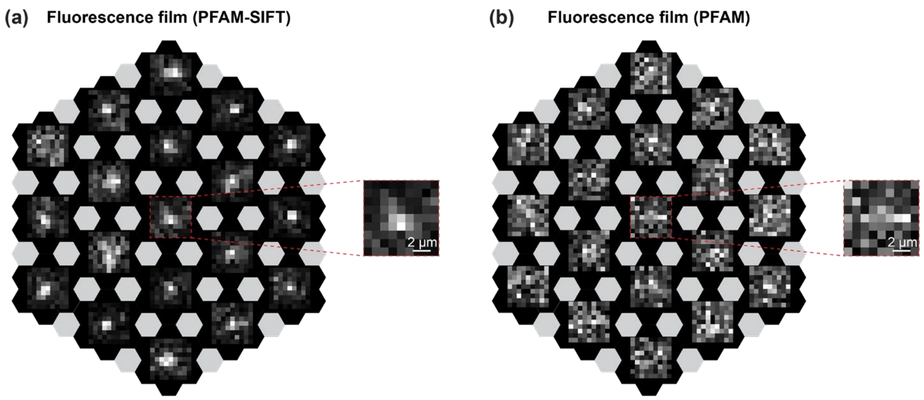


**Supplementary Figure 6. Tip/tilt interference maps of fluorescent film measured using PFAM-SIFT and PFAM.** Same fluorescent film as in ***Fig. 4b***. (**a**) Tip/tilt interference maps for PFAM-SIFT. (**b**) Tip/tilt interference maps for PFAM. Insets: Example interference map (scaled to the sample plane) of a macro-segment.

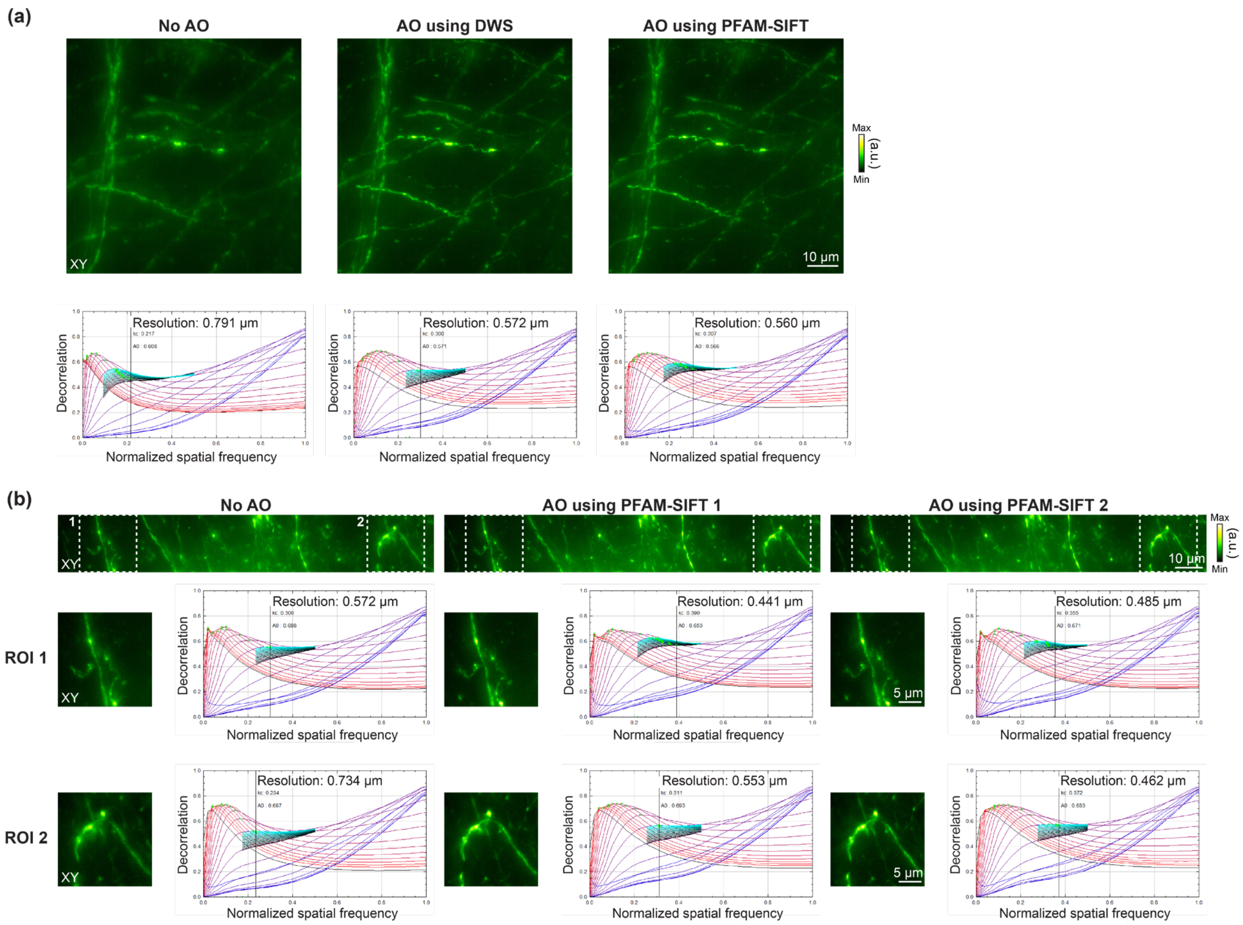


**Supplementary Figure 7. PFAM-SIFT improves lateral resolution of Thy1-GFP line M mouse brain slice images as indicated by Fourier Ring Correlation analysis.** (**a**) (Top) XY Maximum intensity projections (MIPs) from **Fig. 5a**; (Bottom) Fourier Ring Correlation (FRC) plots with estimated resolution values. (**b**) (Top) XY MIPs from **Fig. 5e**; (Middle) XY MIPs of ROI 1 (white dashed box from top row) and corresponding FRC plots; (Bottom) MIPs of ROI 2 (white dashed box from top row) and corresponding FRC plots.

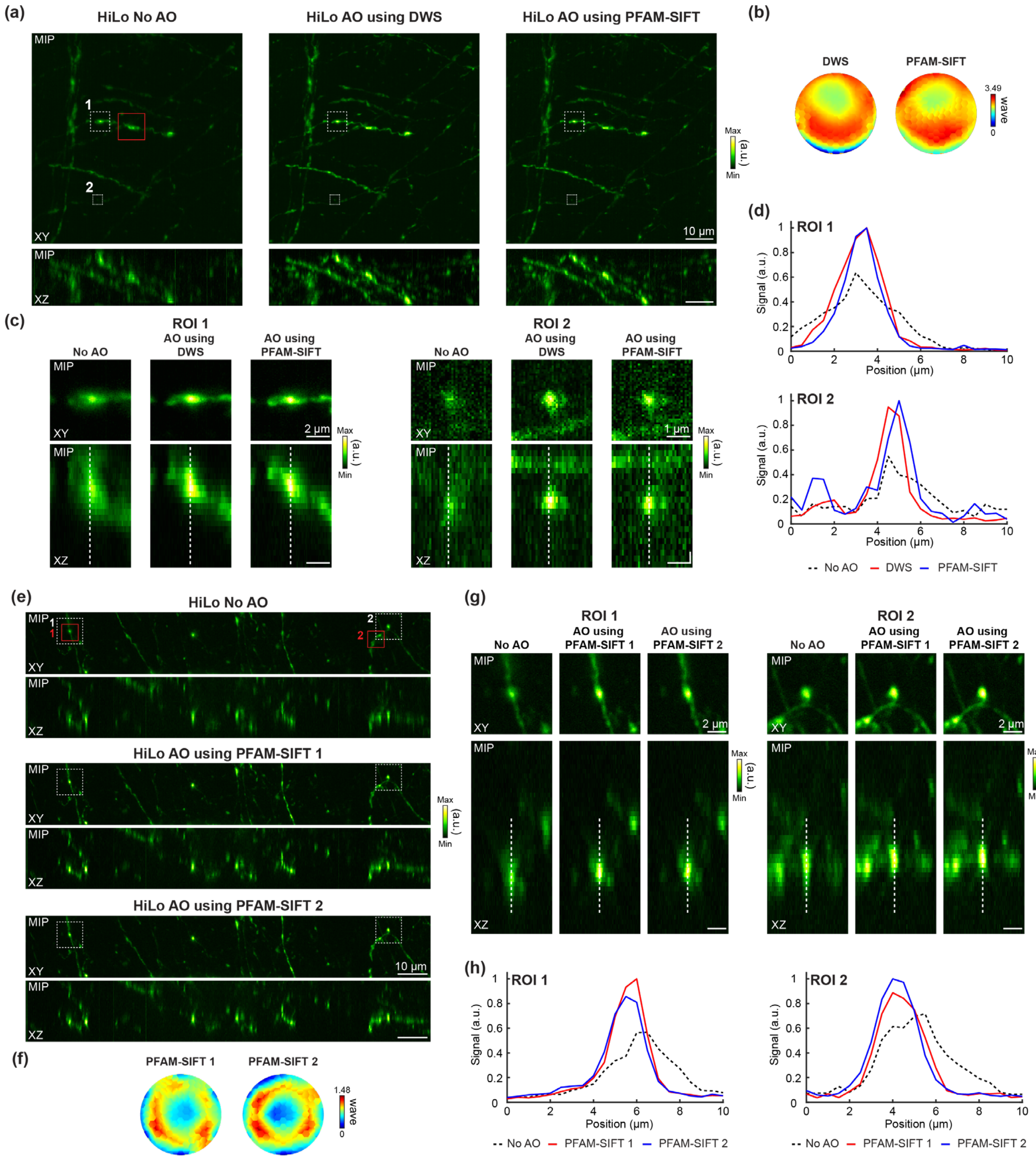


**Supplementary Figure 8. PFAM-SIFT improves HiLo microscopy imaging of Thy1-GFP mouse brain slice.** Same data as in **Fig. 5** but for HiLo images reconstructed from the SI and widefield images using an ImageJ plugin developed by the laboratory of Jerome Mertz (http://biomicroscopy.bu.edu/).

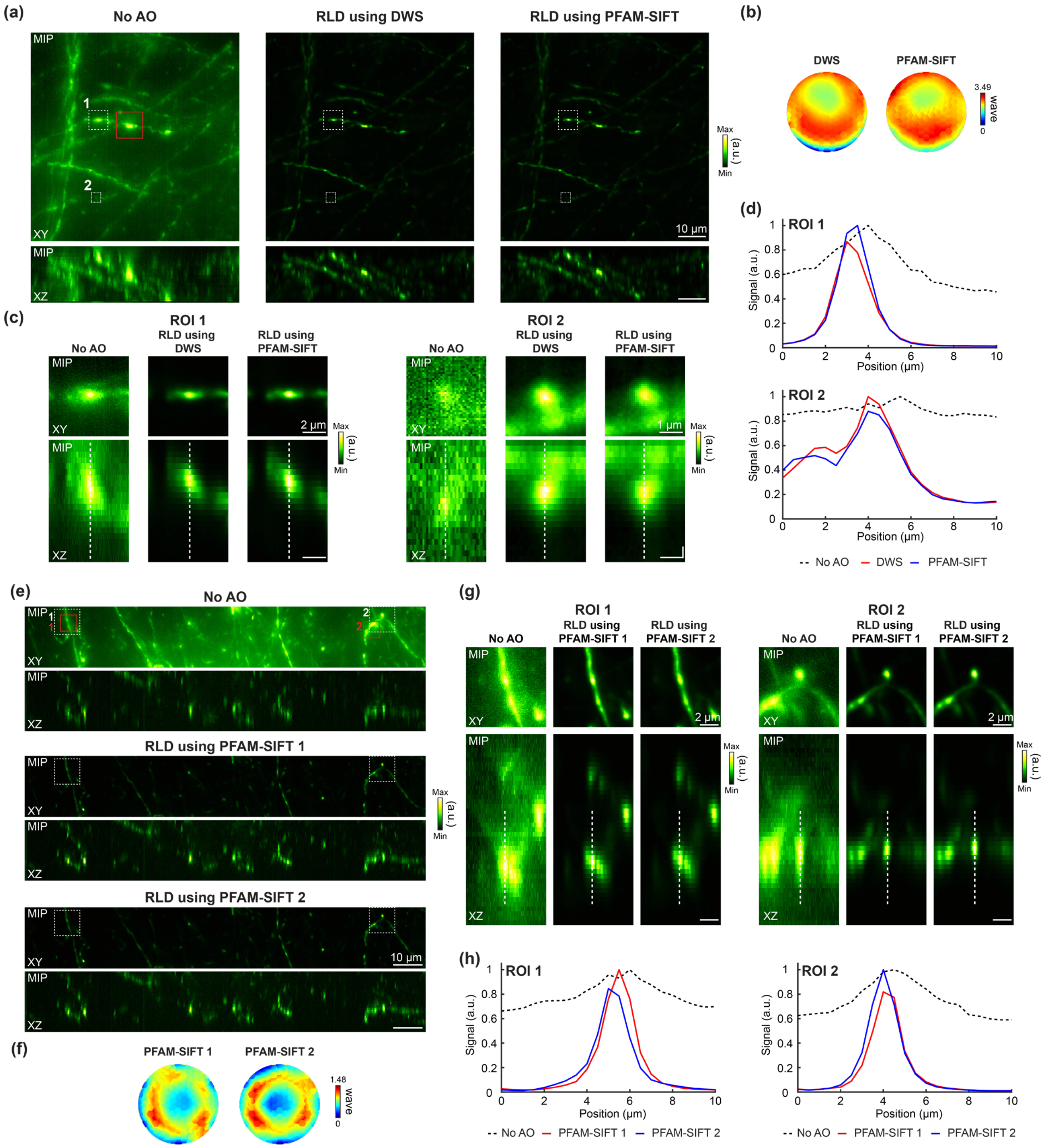


**Supplementary Figure 9. PFAM-SIFT improves deconvolved images of Thy1-GFP mouse brain slice.** Same data as in **Fig. 5** but for images deconvolved with Richardson–Lucy deconvolution (RLD). 'No AO' images are normalized to their own minimum and maximum pixel values, whereas the deconvolved images share calibration bars with the same minimal and maximal pixel values across both conditions: RLD using DWS and PFAM-SIFT (in (**a**), (**c**)), RLD using PFAM-SIFT 1 and PFAM-SIFT 2 (in (**e**), (**g**)).

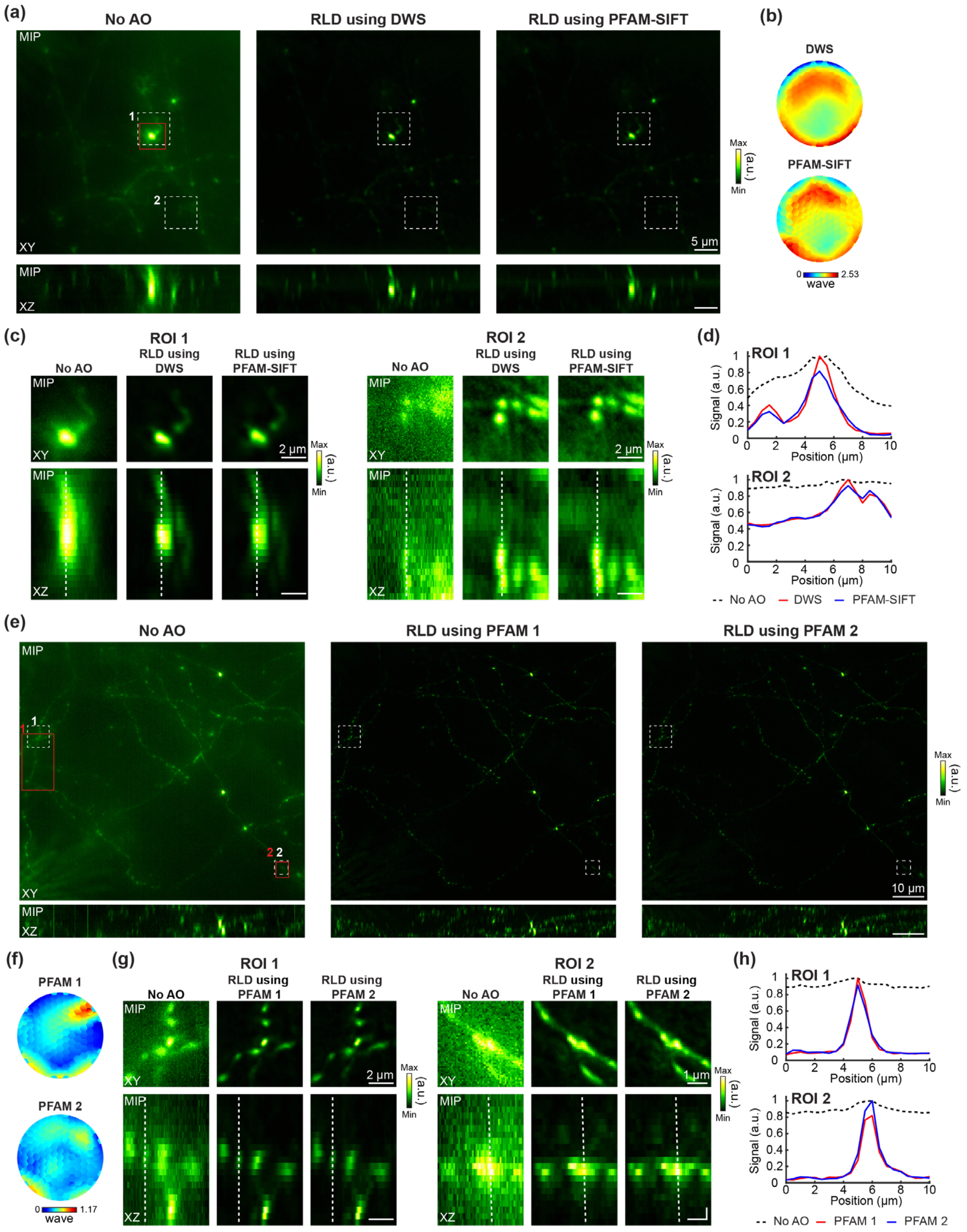


**Supplementary Figure 10. PFAM-SIFT and PFAM improve deconvolved images of tdTomato-expressing neurons in the mouse brain in vivo.** Same data as in **Fig. 6** but for images deconvolved with Richardson–Lucy deconvolution (RLD). 'No AO' images are normalized to their own minimum and maximum pixel values, whereas the deconvolved images share calibration bars with the same minimal and maximal pixel values across both conditions: RLD using DWS and PFAM-SIFT (in (**a**), (**c**)), RLD using PFAM 1 and PFAM 2 (in (**e**), (**g**)).

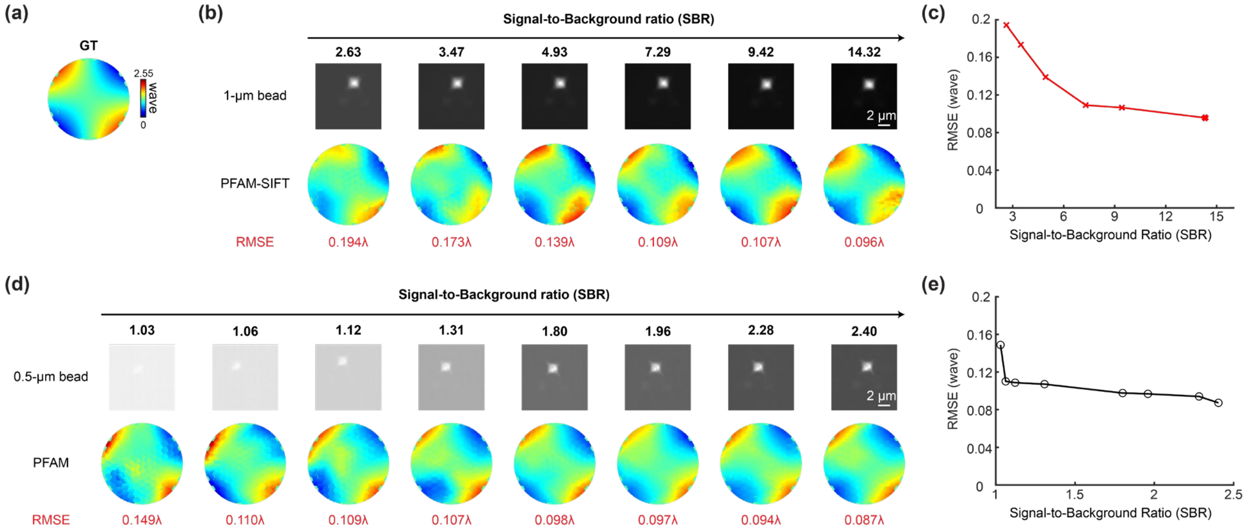


**Supplementary Figure 11. Signal-to-background ratio (SBR) analysis for PFAM-SIFT and PFAM.** (**a**) Ground-truth corrective wavefront for astigmatism of 0.37 waves RMS applied to DM. (**b**) (Top) XY images of a 1-µm-diameter bead under varying SBR conditions were used for PFAM-SIFT and (Bottom) the corrective wavefronts. PFAM-SIFT was iterated once. RMSE: RMS errors between GT and measured wavefronts. (**c**) RMSE vs. SBR in (**b**). (**d**, **e**) Same as (**b**) and (**c**), but with images of a 0.5-µm-diameter bead used for PFAM. PFAM was iterated once. SBR calculation: Otsu's thresholding method was used to segment the image into object and background regions. Mean pixel values of the object and background were then computed, and their ratio was taken as SBR.
